# GenNet framework: interpretable neural networks for phenotype prediction

**DOI:** 10.1101/2020.06.19.159152

**Authors:** Arno van Hilten, Steven A. Kushner, Manfred Kayser, M. Arfan Ikram, Hieab H.H. Adams, Caroline C.W. Klaver, Wiro J. Niessen, Gennady V. Roshchupkin

## Abstract

Deep learning is rarely used in population genomics because of the computational burden and challenges in interpreting neural networks. Here, we propose GenNet, a novel open-source deep learning framework for predicting phenotypes from genetic variants. In this framework, interpretable and memory-efficient neural network architectures are constructed by embedding biological knowledge from public databases, resulting in neural networks that contain only biological plausible connections.

We applied the framework to seventeen phenotypes from a case-control study, a population-based study and the UK Biobank. Interpreting the networks revealed well-replicated genes such as *HERC2* and *OCA2* for hair and eye color and novel genes such as *ZNF773* and *PCNT* for schizophrenia. Additionally, the framework obtained an AUC of 0.74 in the held-out test set and identified ubiquitin mediated proteolysis, endocrine system and viral infectious diseases as most predictive biological pathways for schizophrenia.

GenNet is a freely available, end-to-end deep learning framework that allows researchers to develop and use interpretable neural networks to obtain novel insights into the genetic architecture of complex traits and diseases.

## Introduction

While genome-wide association studies (GWAS) have identified numerous genomic loci associated with complex traits and diseases, the biological interpretation of the underlying mechanisms often remain unclear. Recent GWAS studies with increasingly large sample sizes are resulting greater numbers of significant associations, at an increasing number of independent loci. To illustrate, the latest GWAS for body height based on 700,000 individuals identified more than 3000 near-independent significantly associated single nucleotide polymorphisms (SNPs)^1^. Uncovering a clear biological interpretation from all this information is a challenging task, in which causal variants, genes and pathways need to be identified. In response, many methods such as MAGMA^2^, ALIGATOR^3^ and INRICH^4^, have been developed to obtain a biological interpretation from GWAS summary statistics, providing insights into relevant genes and pathways for defined phenotypes of interest. These methods explore GWAS summary statics and utilize knowledge from annotated biological databases such as NCBI RefSeq^5^, KEGG^6^, Reactome^7^ and GTEx^8^, that have proven to contain crucial information for understanding the underlying biological mechanisms of the human genome.^9^

Given the increasing amount of data available via biobanks and new developments to integrate data, it is now feasible to analyze raw data with more advanced methods. Deep learning is the state of the art in many domains such as medical image analysis^10^ and natural language processing^11^ because of its flexibility and modeling capabilities. In many cases, deep learning yields better performance than traditional approaches, since it scales very well with data size and can model highly non-linear relationships. However, a limitation to deep learning is that these algorithms are often uninterpretable because of their complexity^12,13^. Additionally, genetic data does not lend itself well to the convolution operation, the main driver of the success of deep learning in the imaging domain. Classical fully connected neural networks are not feasible because of the millions of input variants, a fully connected layer would require an infeasible amount of computational time and memory.

To overcome these limitations, we propose GenNet, a novel framework for predicting phenotype from genotype. Within the GenNet framework, biological information from annotated biological sources such as NCBI RefSeq, KEGG and single RNA gene expression datasets, is used to define biologically plausible connections. As a result, neural networks based on this framework are memory efficient, interpretable and yield biological interpretations for their predictions. GenNet is an end-to-end deep learning framework available as a command line tool (https://github.com/ArnovanHilten/GenNet/).

## Methods

The main concept of the GenNet framework is summarized graphically in Figure 1. In this framework, prior knowledge is used to create groups of connected nodes to reduce the number of learnable parameters in comparison to a fully connected neural network. For example, in the first layer, biological knowledge in the form of gene annotations, is used to group millions of SNPs and to connect those SNPs to their corresponding genes. The resulting layer retains only meaningful connections, significantly reducing the total number of parameters compared to a classical fully connected layer. Because of this memory-efficient approach networks in the GenNet framework are able to handle millions of inputs for genotype-to-phenotype prediction. Biological knowledge is thus used to define only meaningful connections, shaping the architecture of the neural network. Interpretability is inherent to the neural network’s architecture; each node is uniquely defined by its connections and represent a biological entity (e.g., gene, pathway). For example, a network that connects SNPs-to-genes and genes-to-output. The learned weights of the connections between layers represent the effect of the SNP on the gene or the effect of the gene on the output. In the network, all neurons represent biological entities and weights model the effects between these entities, together forming a biologically interpretable neural network.

**Figure 1.**
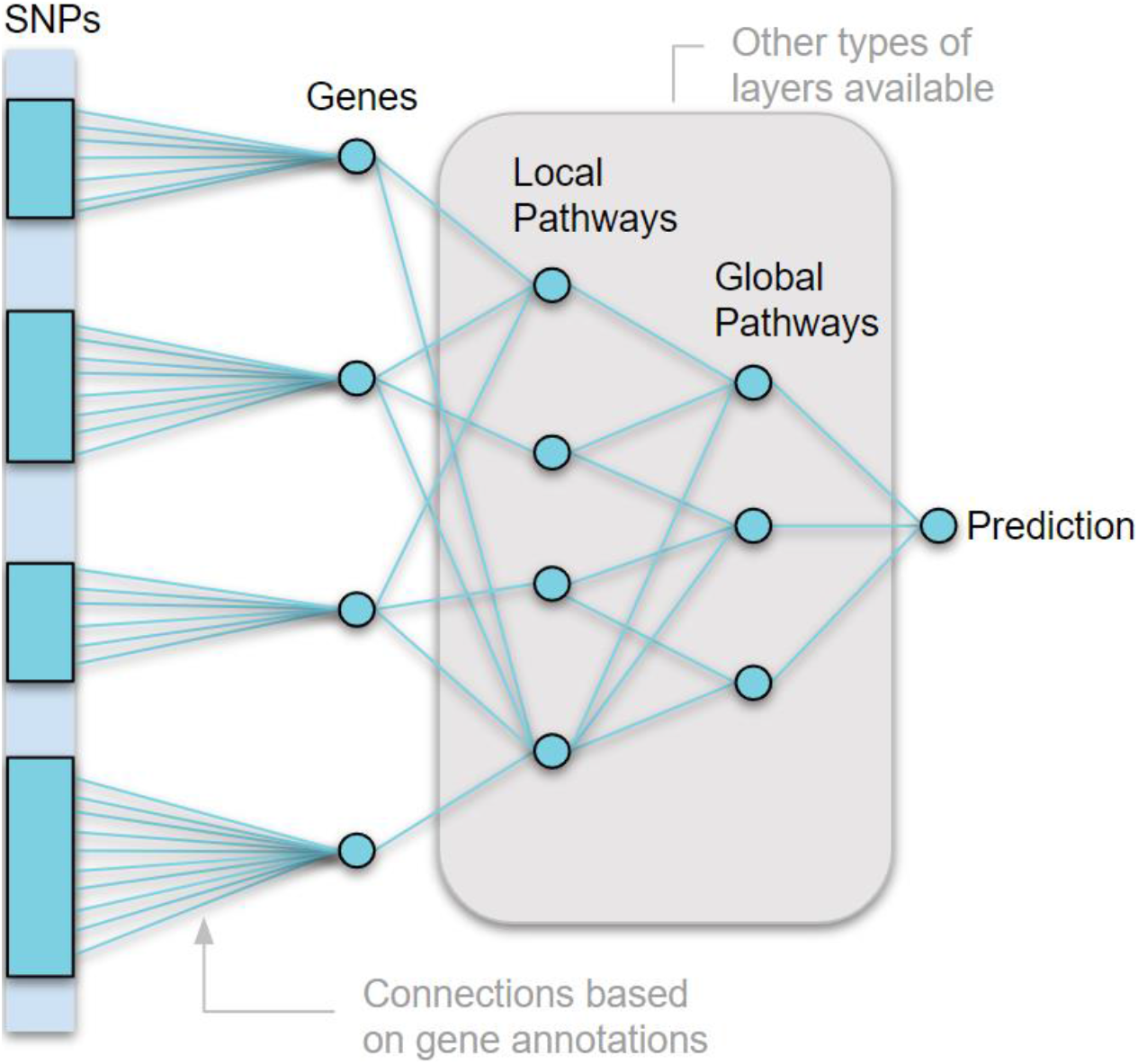
Overview of the GenNet Framework. Neural networks are created by using prior biological knowledge to define connections between layers (i.e., SNPs are connected to their corresponding genes by using gene annotations and genes are connected to their corresponding pathway by using pathway annotations). Prior knowledge is thus used to define each connection, creating interpretable networks.

Many types of layers can be created using this principle. Apart from gene annotations, our framework provides layers built from exon annotations, pathway annotations, chromosome annotations, and cell and tissue type expressions. All these layers can be used like building blocks to form new neural network architectures.

## Results

We built proof of concept simulations as shown in Figure 2A and described in Supplementary Material 1. These simulations demonstrate that the proposed network architecture is interpretable, the strongest weights are assigned to causal variants and genes. Next, we designed experiments to test the network architecture’s performance under a variety of conditions. Performance is dependent on phenotype, neural network architecture and dataset size. To disentangle this, we created simulated data with varying levels of heritability, number of training samples and polygenicity (Supplementary Materials 1). Figures 2B and 2C demonstrate the major trends observed in the simulations. As expected, the network performs best for traits with high heritability, high number of training samples, and low polygenicity.

**Figure 2.**
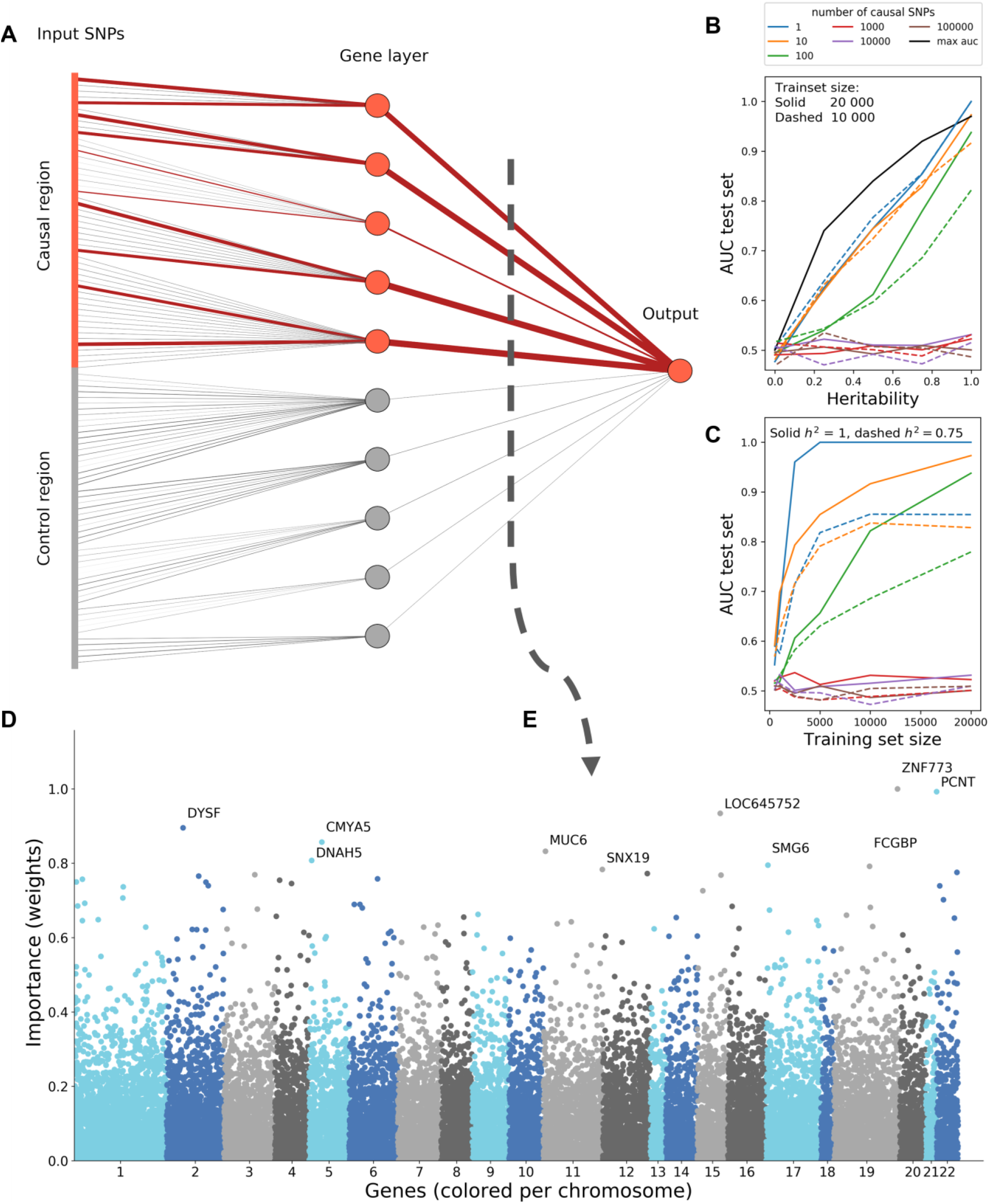
**A)** A simple, non-linear proof of concept. In this simulation, each gene in the causal region has two causal SNPs that cause the simulated disease. The magnitude of the learned weight is represented by the line thickness (contributing causal connections in red, non-contributing control connections in grey). **B)** A secondary set of simulations show the performance of GenNet, expressed as area under the curve, for increasing levels of heritability and training set size **(C)**. The black curve presents the theoretical maximum of the AUC versus heritability. **D)** Manhattan plot showing genes and their relative importance according to the network, here we have shown the results for distinguishing between schizophrenia cases and controls in the Sweden exome study. **E)** This Manhattan plot is a cross section between the gene layer (21,390 nodes) and the outcome of a trained network with 1,288,701 input variants.

Motivated by the proof-of-concept and outcomes of the simulations, we applied the framework to data from multiple sources, including population-based data from the UK Biobank study^14^, the Rotterdam study^15^, and the Swedish Schizophrenia Exome Sequencing study^16^. Phenotypes vary from traits that can be predicted well from only a dozen variants (eye color) to disorders in which thousands of variants explain only a small portion of the variance (schizophrenia and bipolar disorder)^17,18^. The datasets also vary in size and type of data. We used 113,241 exonic variants from imputed microarray-based GWAS data from the Rotterdam study for predicting eye color while we predict fourteen phenotypes in the UK Biobank using 6,986,636 input variants from whole exome sequencing (WES) data. Finally, we used 1,288,701 WES input variants for predicting schizophrenia in the Swedish study. An overview of the main results for networks embedded with gene and pathway information can be found in Table 1. The results for all the experiments, including expression-based networks, can be found in Supplementary 2.

**Table 1.**
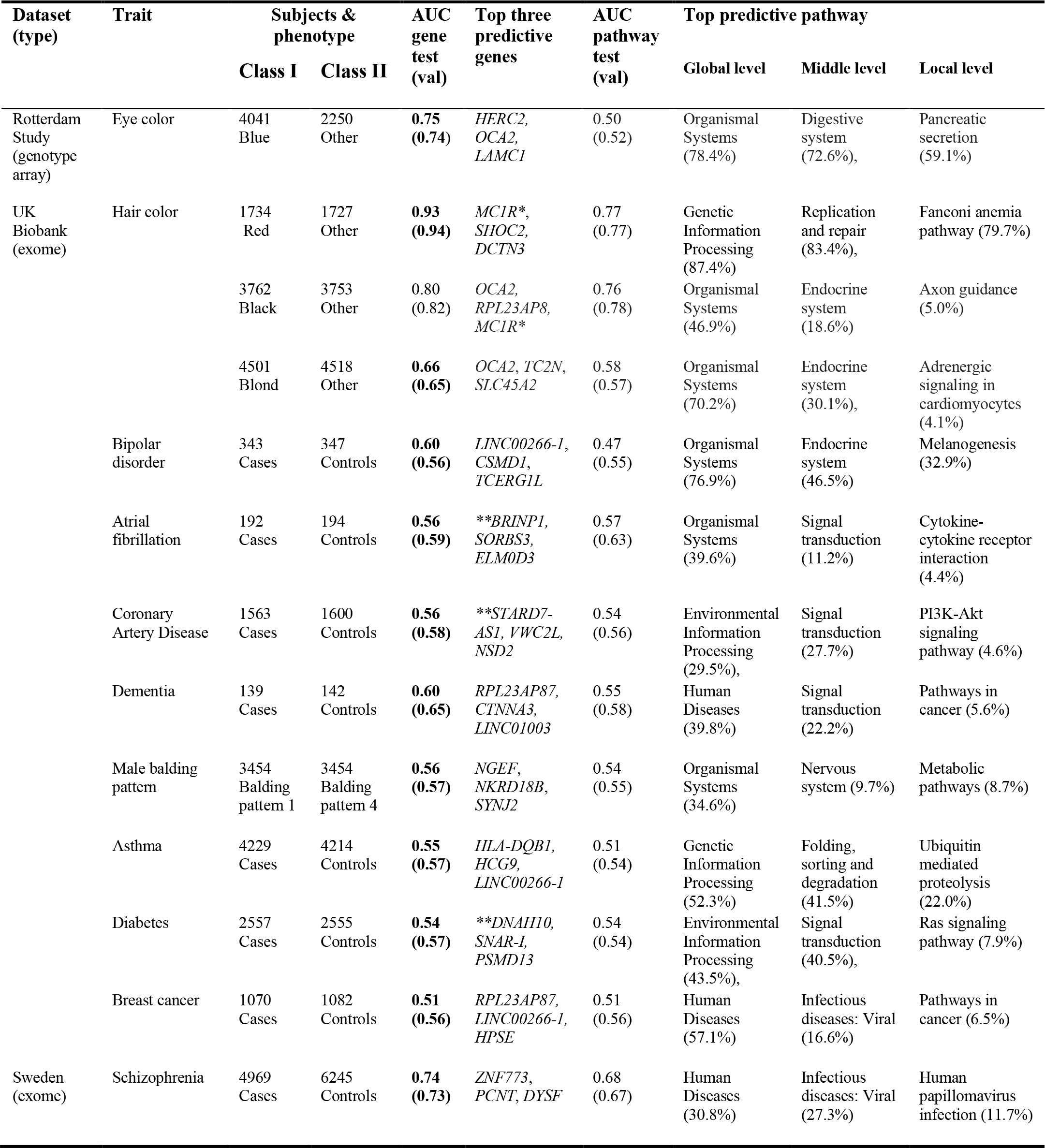
Overview of the main results of the experiments. The performance in AUC for the network with gene-annotations is bold if the network outperformed or matched LASSO regression. Manhattan plots for the genes can be found in Supplementary Materials 3, 4 and 5. *MC1R was not annotated but was identified by linkage disequilibrium. **Many genes contributed to the prediction without clear separation between genes (see Supplementary Materials 4).

Phenotypes with more training samples and that require less variants to obtain high predictive performance, such as eye and hair color, yielded the best performance. Nonetheless, we achieved a good predictive performance for schizophrenia, a highly polygenic disorder, with an area under the curve (AUC) of 0.74 in the held-out test set. All models based on gene annotations outperformed or matched the baseline LASSO logistic regression model with the exception of the black hair color phenotype.

Inspecting the networks, we found that the *OCA2* gene was the most important gene to predict blond, dark and light brown hair color. *OCA2* is involved in the transport of tyrosine, a precursor of melanin.^19^ The *OCA2* signal is likely amplified by the nearby gene *HERC2*, previously identified via functional genetic studies as harboring a strong, long-distance enhancer regulating *OCA2* gene expression underlaying pigmentation variation.^19^ According to the network, *OCA2* and *HERC2* are the two most predictive genes for predicting blue iris color. Both genes were previously identified through GWAS studies of hair and eye color.^20–22^

In the experiments with schizophrenia as outcome, the network was able to classify cases and controls with a maximum accuracy of 0.685 (mean of 0.663 ± 0.014 over 10 runs). Based on all genetic aspects, we estimate the theoretical upper limit for classification accuracy to be roughly 0.72 for schizophrenia (Supplementary Materials 7). The model achieved a maximum area under the receiver operating curve of 0.738 (mean of 0.715 ± 0.016) in the held-out test set over 10 runs, thereby considerably outperforming the LASSO logistic regression baseline which obtained a maximum AUC of 0.649 (mean of 0.644 ± 0.003). The genes *ZNF773, PCNT* and *DYSF* were assigned the highest weights by the neural network and were thus considered to be the most predictive genes for schizophrenia. However, the polygenic nature of schizophrenia can be seen when comparing the Manhattan plot of the genes (Figure 2D) to other phenotypes in this study (Supplementary Materials 3,4 and 5). More genes contributed to the prediction for schizophrenia than for the other phenotypes examined.

Figure 3 shows the relative importance of the pathways for predicting schizophrenia. The neural network, embedded with the KEGG pathway information, obtained an AUC of 0.68 in the test set. Human diseases (30.8%), organismal systems (26.7%) and genetic information processing (26.5%) were the main contributors to the neural network’s prediction of schizophrenia. The contribution of human diseases was mainly driven by viral infectious diseases (27.3%) which can be subdivided further into human papillomavirus infection (11.7%), herpes simplex infections (3.7%), and human cytomegalovirus infection (3.2%), as well as genes and SNPs assigned to these diseases. The ubiquitin mediated proteolysis pathway, a subset of the genetic information pathway, had a relative importance of 10.0%.

**Figure 3.**
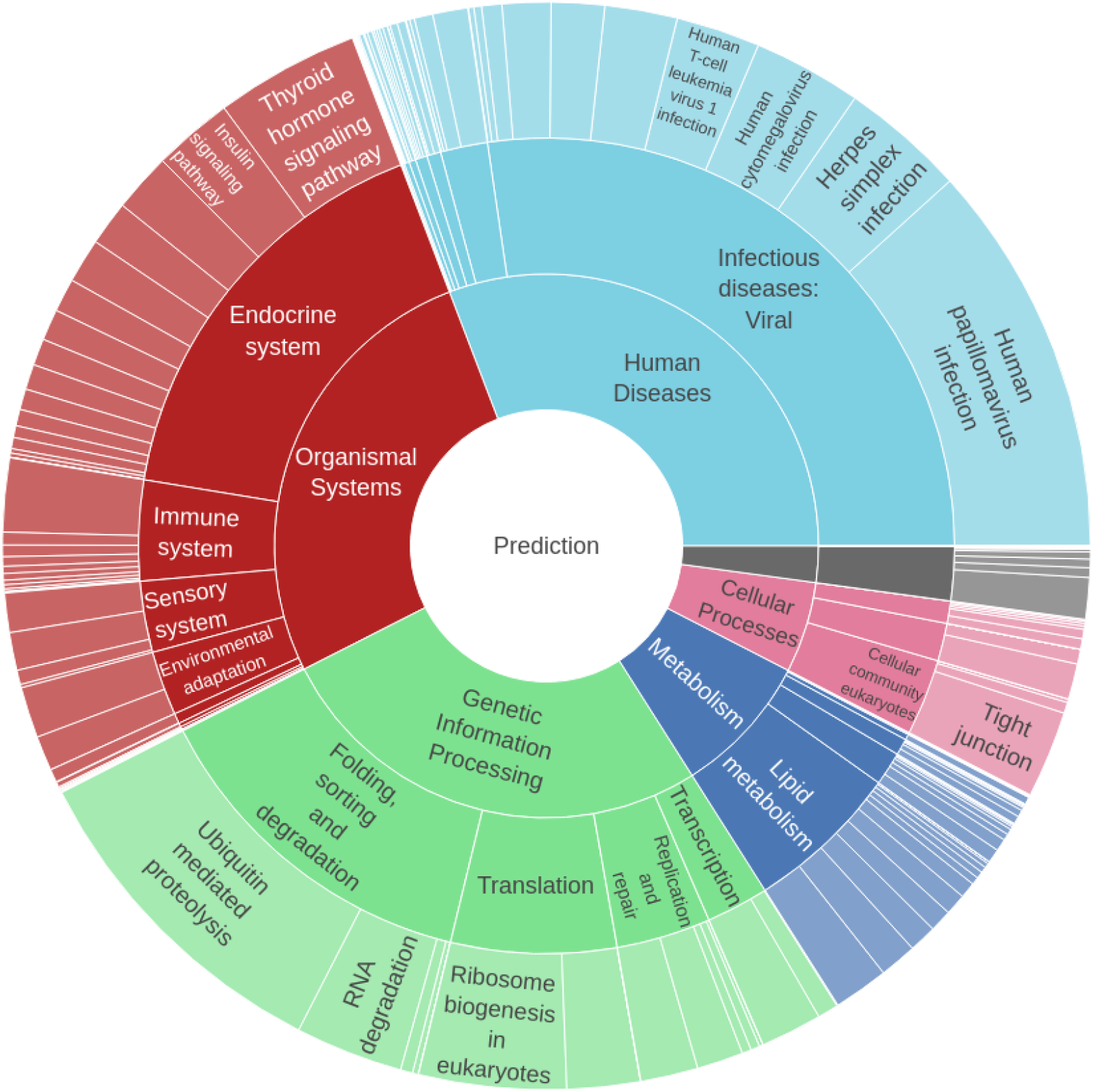
Sunburst plot of the important pathways for predicting schizophrenia. A neural network with layers based on KEGG pathway information was trained to predict schizophrenia. The relative importance was calculated for each pathway using the learned weights of this neural network. The sunburst plot is read from the center, which shows the output node with the prediction of the neural network. The inner ring represents the last layer with the highest-level pathways, followed by the mid-level pathways and finally the lowest-level pathways. The gene and SNP layer are omitted for clarity.

To investigate if the most predictive genes are concentrated in a single cell or tissue type, we used three different gene expression datasets processed by Finucane et al. (2018)^23^ to create layers subsequent to the gene-based layer. The relative importance of the trained networks showed a widespread signal with small contributions by many cell and tissue types. When GTEx data was embedded in the network, uterine genes had the greatest relative importance (4.2%). For GTEx brain expression data the frontal cortex (15.1%) had the strongest contribution, whereas for immune cell types mesenteric lymph nodes (1.23%) contributed most. An overview of the experiments performed, including GTEx and other expression-based networks for other phenotypes, can be found in Supplementary Material 2.

## Discussion

We presented a novel framework for predicting phenotypes from genotype with interpretable neural networks. The proposed neural networks have connections defined by prior biological knowledge only, reducing the number of connections and therefore the number of trainable parameters. Consequently, the networks can be trained on a single GPU and offer a biological interpretation for their predictions.

In the first set of experiments, simulations showed the network’s performance when varying the degree of heritability, polygenicity and sample size. In the second set of experiments the framework was applied to UK Biobank, Rotterdam study and Swedish Schizophrenia Exome Sequencing Study data. In these experiments, phenotypes with widely different heritability, sample size and polygenicity were predicted with generally high predictive performance. For example, we obtained an AUC of 0.96 for red hair color, an AUC of 0.74 for blue eye color and an AUC of 0.74 for predicting schizophrenia. For schizophrenia, polygenic risk scores from genome wide association studies obtain similar AUCs, ranging from 0.49 to 0.85, while spanning the whole genome and using larger sample sizes.^17^

Interpretation of the trained networks revealed well-replicated genes for traits with a known etiology such as *HERC2* and *OCA2* for eye color and *OCA2* and *TC2N* for hair color.^20–22,24^ For schizophrenia, a disorder with an unclear etiology, the network identified novel genes, including *ZNF773* and *PCNT*. In previous studies, *PCNT* was only nominally associated with schizophrenia, although it is known to have interactions with *DISC1* (Disrupted in Schizophrenia-1).^25^ There is a strong correlation between (prenatal) viral infections and increased risk for developing schizophrenia.^26^ It is therefore interesting that by using the KEGG pathway information to create layers, we identified the viral infectious diseases pathway with its associated pathways, genes and SNPs to be the most important for predicting schizophrenia. The results of these experiments reveal that with GenNet framework can provide insights across multiple functional levels, identifying which SNPs, genes, pathways and tissue types are important for prediction.

In this study, WES data and exonic variants from microarrays have been used, however the principles in the GenNet framework can be extended to handle diverse types of input, including genotype, gene expression and methylation data or combinations thereof. Similarly, we explored only a fraction of the possible layers as building blocks for our networks. Any data that create unique biological groups can be used in the framework to create new layers. However, the quality of the resulting network is highly dependent on the quality of the data used. For example, networks built with biological pathway data performed generally worse than networks using only gene annotations since many genes are not annotated to pathways. But with new and better data from projects such as GTEx^8^, ENCODE ^27^ and KEGG^6^ the quality of the layers will only improve. Moreover, given the increasing sample sizes of biobanks in coming years and the development of algorithms, which allow distributed deep learning between cohorts without sharing raw data^28^, we foresee that our framework can be easily adopted for such settings and can be widely used for quantitate traits and functional mapping analysis.

In conclusion, we developed a freely-available framework, which can be used to build interpretable neural networks for genotype data by incorporating prior biological knowledge. In addition to computationally efficient, the architectures are interpretable, thereby alleviating one of the most important shortcomings of neural networks. We have demonstrated the effectiveness of this novel framework across multiple datasets and for multiple phenotypes. Given that each network node is interpretable, we anticipate this approach to have wide applicability for uncovering novel insights into the genetic architecture of complex traits and diseases.

## Online Methods

### Sweden Schizophrenia Exome Sequencing Study

Sweden-Schizophrenia Population-Based Case-Control Exome Sequencing study (dbGaP phs000473.v2.p2), is a case control study with 4969 cases and 6245 controls.^16^ All individuals aged 18-65, have parents born in Sweden, and provided written informed consent. The following inclusion criteria were used for cases: at least two times hospitalized with schizophrenia discharge diagnosis; no hospital register diagnosis consistent with a medical or other psychiatric disorder that mitigates the schizophrenia diagnosis. Cases were excluded if they had a relationship closer than 2nd degree relative with any other case. Controls were matched to cases and controls were never hospitalized with a discharge diagnosis of schizophrenia. Controls were excluded if they had any relation to either case or control.

The downloaded plink files were converted using HASE^29^ to hierarchical data format (HDF), a format that allows fast and parallel data reading. After conversion, the data was transposed and SNPs without any variance were removed (1,288,701 SNPs remain). The data was split in a training, validation and test set (ratio of 60/20/20), while preserving the ratio cases and controls.

### UK Biobank

We applied the framework to multiple phenotypes in the UK Biobank using the first release of the WES data, providing whole exome sequencing for 50,000 UK Biobank participants.^30^ Phenotypes are self-reported using touchscreen questions in the UK Biobank Assessment Centre. Similar to the Sweden cohort all variants without variance were removed, data was converted to hierarchical data format, and transposed. For every phenotype an equal number of cases and controls were sampled. The resulting dataset is split in a train, validation and a test set (ratio of 60/20/20). Related cases, and cases with related controls, (kinship > 0.0625) are all in the training set. This is done under the assumption that related cases and controls could ease training, the shared genetic information could steer the network towards the discriminatory features. The validation and test sets contain only unrelated cases and controls within and between sets. Unrelated controls are randomly sampled and added to gain an even distribution between cases and controls in all sets. Misaligned SNPs and sex chromosomes were masked in the first layer and therefore not included in the study.

### Rotterdam Study

The Rotterdam study is a prospective population-based cohort study.^15^ We used the first cohort consisting of 6291 participants, genotyped using the Illumina 550K and 550K duo arrays. Samples with low call rate (<97.5%), excess autosomal heterozygosity (>0.336) or sex-mismatch were excluded, as were outliers identified by the identity-by-state clustering analysis (outliers were defined as being >3 standard deviation (SD) from population mean or having identity-by-state probabilities >97%). For imputation the Markov Chain Haplotyping (MACH) package version 1.0 software (Imputed to plus strand of NCBI build 37, 1000 Genomes phase I version 3) and minimac version 2012.8.6 were used (call rate >98%, MAF >0.001 and Hardy–Weinberg equilibrium P-value > 10^−6^). From here on, processing steps are identical as described for Sweden Schizophrenia resulting in a dataset with 113,241 exonic variants for 6291 subjects. For each subject, both eyes were examined by an ophthalmological medical researcher and eye (iris) color was categorized into three categories; blue, intermediate and brown using standard images and based on the predominant color and pigmentation.^31^

### Prior knowledge

All SNPs were annotated using Annovar^32^. From these annotations a sparse connectivity matrix was created, connecting the SNPs to their corresponding genes. Connectivity matrices between SNPs, exons and genes were made using intron-exon annotations. A mapping between genes and pathways was made using GeneSCF^33^ and the KEGG database^6^. GTEx tissue-expression masks were made using the fully processed, filtered and normalized gene expression matrices for each tissue directly obtainable from the GTEx website^8^. All expression based layers in this paper are created from derived t-score statistics^23^ to ensure that nodes are uniquely defined by their connections and therefore interpretable. There are many possibilities to expand the GenNet framework, it can create layers from any data that groups data uniquely. For example the framework contains layers based on single cell expression is created from data released by FUMA^34^.

### Neural network architecture

In the GenNet framework, layers are available built from biological knowledge such as; exon annotations, gene annotations, pathway annotations, cell expression and tissue expression. Information from these resources is used to define only meaningful connections, shaping an interpretable and lightweight neural network, allowing evaluation of millions of input variants together. These networks bear similarities to the first generation of neural networks and recently interest for these networks has rekindled for biological applications.^35–38^

The GenNet architecture aims to reduce the dimensionality of the network with a minimal loss of information. By using layers based on prior biological knowledge, the network is reduced in a biologically sensible way. During training, weights in the network are optimized to predict the outcome. The network learns the relation between the SNPs, genes and other biological information used to create the layers. Because the input to each layer is normalized, the important connections for predicting the phenotype will receive higher absolute weights while unimportant connections will receive lower absolute weights.

All models were trained on a single GPU with 11 GB memory (Nvidia GeForce GTX 1080 Ti) and converged within five days. During the experiments, we noticed it is beneficial to add an L1 penalty on the weights in the last dense layer, similar to LASSO logistic regression. Regularization constraints the search space which leads to better generalization and interpretation.

### Interpretation

Interpretation of the network is straightforward due to the simplicity of the concept, the stronger the weight is the more it contributes to the final prediction of the network. The simplest network in the framework, a network built by gene annotations, can be seen as ~20 000 (number of genes) parallel regressions followed by a single logistic regression. The learned weights in these regressions are similar to the coefficients in logistic regression. Especially the last node, a single neuron with a sigmoid activation, 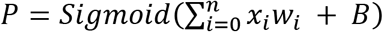 is very similar to logistic regression 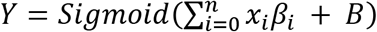.

The inputs in logistic regression are generally normalized to compare coefficients (*β*). In the neural network this is achieved by batch normalization (without center and scaling), normalizing the weights (*w*) after every activation. The weights can then be compared and used as an importance measure. Since batch normalization is a batch-wise approximation, the learned weights can be multiplied with the standard deviation of the activations for a more accurate estimate. For Manhattan plots, the normalized, absolute weights between the gene layer and the output is used.

In larger networks, the relative importance is calculated by multiplying the weights for each possible path from the end node to the input SNPs. At each input we obtain then a value denoting its contribution. These values are then aggregated (summed) according to the groups of the subsequent layers to get the importance estimate for each node in the network.

It is important to note that in this work (relative) importance is more similar to effect size or odds ratio than to statistical significance. The weights represent the direction and effect a gene or pathway has on the outcome of the network.

### Implementation

Technically, the computational performance of the implemented Keras/Tensorfow^39,40^ layer should be on par or an improvement over similar layers. It is implemented using sparse matrix multiplication, making it faster than the slice-wise locallyconnected1D layer and more memory efficient than dense matrix multiplication. The layer is friendly to use, with only one extra input compared to a normal dense layer. This extra input, the sparse connectivity matrix, is made with prior knowledge and describes how neurons are connected between layers.

The networks behave similar to normal fully connected artificial neural networks but is pruned by removing irrelevant connection 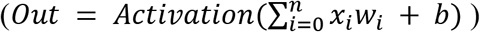 With *w* as a sparse matrix with learnable weights, initialized with a sparse connectivity matrix defining connections. The networks are optimized using the ADAM or Adadelta optimizers, using weighted binary cross entropy with weights depending on the imbalance of the classes, and are all trained on a single GPU. For regression tasks, mean squared error is used as a loss function in combination with ReLu activations (Supplementary Materials 7). The GenNet framework is available as command line tool and includes functionalities to convert data, to create, train and evaluate neural networks and to plot the results.

### Baseline

As a baseline method, LASSO logistic regression was implemented in Tensorflow by using a dense layer of a single neuron with a sigmoid activation function and L1 regularization on weights. Polygenic Risk Scores (PRS) were not used since we used only variants from the exome.

### Upper bound of classification accuracy for predicting phenotypes

In contrast to the application of deep learning for image analysis, where the performance is not limited, in application for genetic analysis we are limited by the information in DNA relevant for trait or disease, i.e., by the heritability. Therefore, we cannot expect the performance of the neural network above certain value.

Population characteristics can be used to calculate the upper bound of performance for a classifier for any trait. This can be done by creating a confusion matrix. The accuracy between true and false positives for a perfect classifier, based solely on genetic inputs, is given by the concordance rate between monozygotic twins. It is impossible to predict better based solely on genetic code than the rate a trait occurs in people with virtually the same genetic code. The chance of misclassifying a control should be better than the prevalence, which is often close to zero for most diseases. Creating a confusion matrix can give insights in the upper bound for accuracy, sensitivity and specificity in the dataset. An example for schizophrenia in our dataset can be found in Supplementary Materials 7.

## Code availability

GenNet is an open-source framework usable from command line. GenNet and its tutorials, including how to build new layers and networks from prior knowledge, can be found on: https://github.com/arnovanhilten/GenNet/

## Data availability

Code to run and generate data for the simulations are publicly available on GitHub. The genetic and phenotypic UK Biobank data are available upon application to the UK Biobank (https://www.ukbiobank.ac.uk/). Access to the Sweden-Schizophrenia Exome Sequencing study can be requested on DBGaP (https://www.ncbi.nlm.nih.gov/gap/) (dbGaP phs000473.v2.p2).

## Acknowledgements

This work was funded by the Dutch Technology Foundation (STW) through the 2005 Simon Steven Meester grant 2015 to W.J. Niessen. This research has been conducted using the UK Biobank Resource under Application Number 23509. This work was carried out on the Dutch national e-infrastructure with the support of SURF Cooperative (application number 17610). The Rotterdam study is supported by the Netherlands Organization for Scientific Research (NWO, 91203014, 175.010.2005.011, 91103012).

## Author contributions

A.H and G.R conceived and designed the method. A.H performed experiments and implemented the method. G.R and W.N supervised the work. M.A.I, C.K, H.A, M.K and S.A.K. provided or gave access to data. A.H, G.R, W.N, M.K., M.A.I., S.A.K, wrote, revised and approved the manuscript.

## Additional Information

Correspondence and requests should be addressed to A.H or G.R.

