## Supplementary Materials for "GenNet framework: interpretable neural networks for phenotype prediction"

#### Table of contents

Supplementary 1. Simulations ... 3

1.1 Proof of concept ... 3

1.2 Synthetic data ... 4

1.3 Genotype data, simulated phenotype ... 5

Supplementary 2. Overview of the experiments ... 6

2.1 Gene networks ... 6

2.2 Pathway networks ... 8

2.2 GTEx expression networks ... 10

2.2 ImmGen expression networks ... 11

Supplementary 3. Sweden-Schizophrenia population-based case-control study ... 12

3.1 Gene network ... 13

3.2 Gene network with an L1 constraint on the weights ... 14

3.3 Pathway network ... 15

3.4 Gene-cell network & gene-GTEx networks ... 16

Supplementary 4. UK Biobank ... 19

4.1 Bipolar disorder ... 19

4.2 Skin color ... 20

4.3 Hair color: one versus all ... 21

4.4 Hair color: one vs one ... 24

4.5 Male balding pattern ... 25

4.6 Breast cancer ... 26

#### Supplementary 1. Simulations

##### 1.1 Proof of concept

This experiment is easily reproduced by running: *https://tinyurl.com/y8hh8rul* online.
The first experiment is a simple experiment designed to show the working principle and how to interpret the results. 200 SNPs are simulated by drawing from a binomial distribution. These SNPs are assigned to 10 genes. The first 5 are set to be causal for our phenotype while the last 5 function as controls. The sizes of the genes are varied to observe the effects of different sizes on the importance, for every causal gene there is a control gene with the same size. (Gene sizes = [50, 30, 10, 5, 5, 50, 30, 10, 5, 5]. The training, validation and test set consist of 10000, 2000 and 2000 subjects respectively (50 % cases vs 50% controls). To ensure equal importance of the genes (even with different gene size) only 1 SNP per gene is set to be causal. Therefore, there are 5 different subtypes of the disease each subtype prevalent in 1/5th of the simulated population.

This is a relatively simple problem and the neural network performs as expected with a near perfect score (AUC of 0.99) in the test set. It can be seen in figure 1 that the network identifies the first five genes to be causal and the last five control genes to be of insignificant importance. Inspection of the weights between SNPs and genes identifies the causal SNPs without any unambiguity for all causal genes. The direction of effect is easily derived by the sign of the weights. Subtypes of the simulated phenotype can be distinguished by activation patterns.

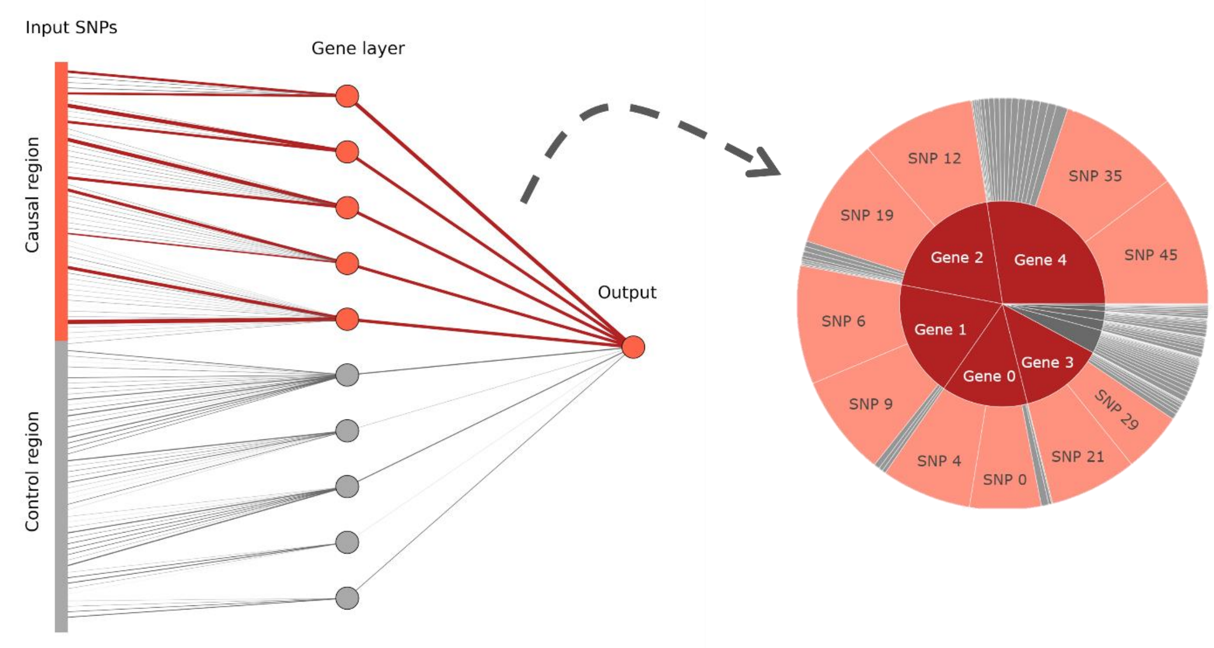

*Supplementary Figure 1: A simple, non-linear proof of concept. In this simulation, each gene in the causal region has two causal SNPs that cause the simulated disease. The magnitude of the learned weight is represented by the line thickness (contributing causal connections in red, non-contributing control connections in grey). The right side shows how this converts to a sunburst plot.*

#### 1.2 Synthetic data

In this simulation the performance of the network is evaluated when heritability, polygenicity and training set sample size are varied. 100,000 SNPs are simulated for N subjects by drawing from a binomial distribution (n=2, p=0.3). Causal SNPs are selected randomly and assigned equal effect size. The number of causal SNPs is controlled by the polygenicity variable. Liability is calculated according to: $Liability = \sqrt[2]{h^2}g + q\sqrt[2]{1-h^2}$ with $h^2$ as heritability, $g$ as genomic risk (effect size multiplied by genotype matrix) and with $q$ as random noise.
Phenotype is based on a liability-threshold model with the threshold half a standard deviation above the mean liability.

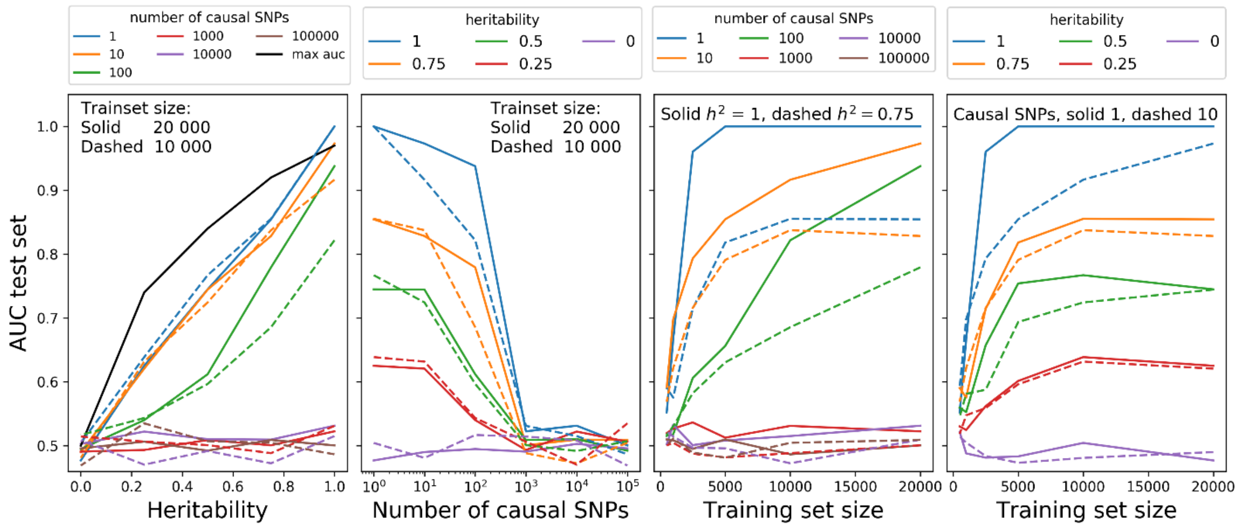

*Supplementary Figure 2: Simulations with synthetic data showing the performance of GenNet expressed in the area under the curve for increasing levels of heritability, training set size and the number of causal SNPs (polygenicity). In total 540 simulations have been run with each 100,000 input variants In black the theoretical maximum of the AUC versus heritability.*[1]

Supplementary figure 3 gives an indication of the expected performance of a 3-layer network for different heritability, polygenicity and sample sizes. The indications are conservative, the network performed better in human genotype than in the simulated data. In genotype data the network obtained better performance for smaller sample sizes and for phenotypes with more causal SNPs than suggested by these simulations. This could be due to the absence of linkage disequilibrium in the simulated data or it could be an artifact of how the phenotype is constructed in the simulations.

##### 1.3 Genotype data, simulated phenotype

The simulations are constructed from real data, the genotype matrix from the schizophrenia Sweden study served as a basis for this simulation. New labels were generated for each patient based on the following pseudo code:

Randomly select a set of **P** genes.
Randomly sample a number of SNPs in each gene (**N**).
Effect of each SNP    Causal    **E** = (1/**P**)/(**N**/2),
    Non-causal    **E** = 0
Phenotype risk = **E** * **M** (Genotype matrix Sweden **M**)
Threshold for cases is the median phenotype risk, in order to get a case control of 50/50.

This experiment was designed to show the effect of LD blocks on the behavior of the neural network. Similar as seen in genome wide association studies, the found genes are not guaranteed to be causal.

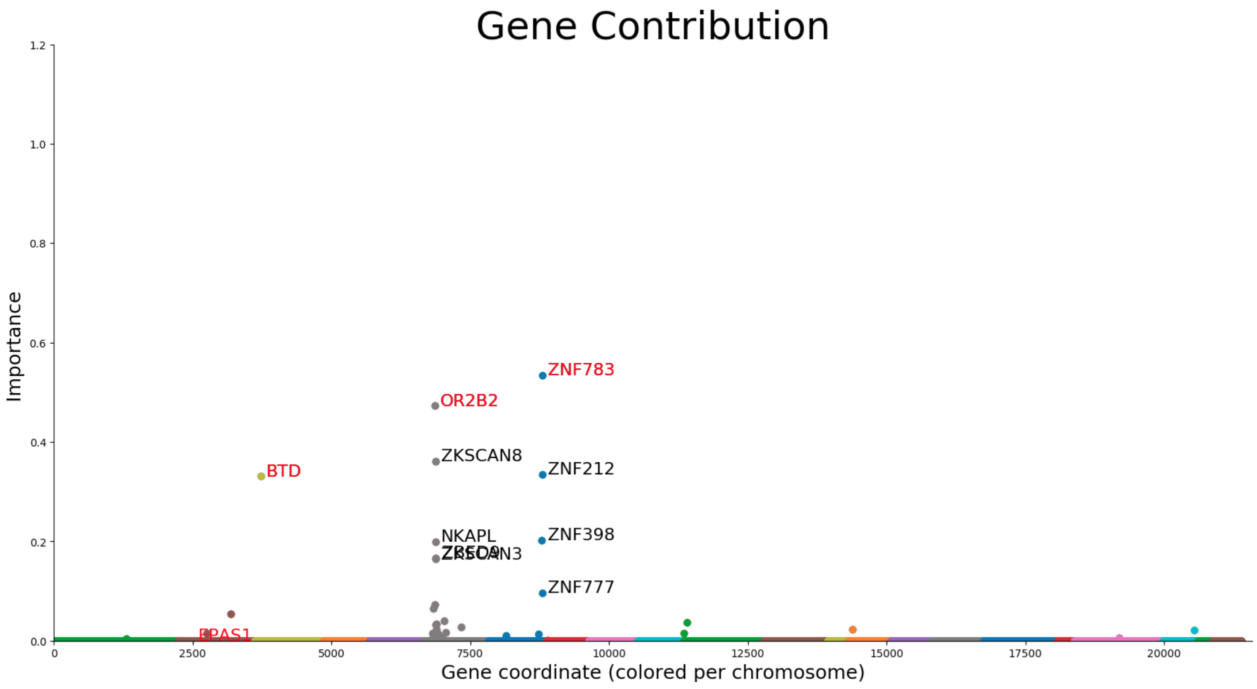

Supplementary Figure 3. Behavior of the Network for LD. Causal genes are in red, non-causal genes are in black. We can observe that the neural network tags SNPs similar to GWAS. The LD structure adds predictive value to non-causal genes due to the linkage disequilibrium.

### Supplementary 2. Overview of the experiments

#### 2.1 Gene networks

| Dataset (type) | Number of input variants | Trait | Subjects & phenotype | | AUC LASSO (val) | AUC LASSO (test) | AUC GenNet (val) | AUC GenNet (test) | GenNet: top three most important genes |
|---|---|---|---|---|---|---|---|---|---|
| | | | Class I | Class II | | | | | |
| Rotterdam (genotype array) | 113,241 (exonic) inputs of 16,628 genes | Eye color | 4041 Blue | 2250 Other | 0.68 | 0.69 | 0.74 | 0.75 | *HERC2, OCA2, LAMC1* |
| UK Biobank (exome) | 6,986,636 input variants of 15,827 genes | Hair color | 4501 Blond | 4518 Other | 0.62 | 0.61 | 0.65 | 0.66 | *OCA2, TC2N, SLC45A2* |
| | | | 15684 Dark brown | 15918 Other | 0.57 | 0.59 | 0.69 | 0.70 | *OCA2, TC2N, SPIRE2* |
| | | | 1734 Red | 1727 Other | 0.67 | 0.70 | 0.94 | 0.93 | *MC1R**, *SHOC2, DCTN3* |
| | | | 16208 Light brown | 16029 Other | 0.61 | 0.62 | 0.62 | 0.62 | *OCA2, TC2N, NOX4* |
| | | | 3762 Black | 3753 Other | 0.85 | 0.82 | 0.83 | 0.81 | *OCA2, RPL23AP87, SPIRE2* |
| | | Skin color | 1883 Fair | 1894 Dark | 0.97 | 0.98 | 0.98 | 0.98 | *RPL23AP87, SMARCAD1, GLP1R* |
| | | Male balding pattern | 3454 Balding pattern 1 | 3454 Balding pattern 4 | 0.57 | 0.57 | 0.56 | 0.57 | *NGEF, NKRD18B, SYNJ2* |
| | | Atrial fibrillation | 192 Cases | 194 Controls | 0.64 | 0.43 | 0.59 | 0.56 | ** *BRINP1, SORBS3, ELM0D3* |
| | | Coronary Artery Disease | 1563 Cases | 1600 Controls | 0.57 | 0.55 | 0.58 | 0.56 | ** *STARD7-AS1, VWC2L, NSD2* |
| | | Bipolar disorder | 343 Cases | 347 Controls | 0.56 | 0.59 | 0.56 | 0.60 | *LINC00266-1, CSMD1, TCERG1L* |
| | | Diabetes | 2557 Cases | 2555 Controls | 0.55 | 0.54 | 0.57 | 0.54 | ***DNAH10, SNAR-I, PSMD13* |
| | | Dementia | 139 Cases | 142 Controls | 0.55 | 0.58 | 0.65 | 0.60 | *RPL23AP87, CTNNA3, LINC01003* |

| | | | | | | | | | |
|---|---|---|---|---|---|---|---|---|---|
| | | Allergies | 10242 Cases | 10187 Controls | 0.51 | 0.51 | 0.53 | 0.52 | *****AC025039.1, AC004052.1, VPS45*** |
| | | Breast cancer | 1070 Cases | 1082 Controls | 0.53 | 0.52 | 0.56 | 0.52 | *RPL23AP87, LINC00266-1, HPSE2* |
| | | Asthma | 4229 Cases | 4214 Controls | 0.53 | 0.51 | 0.57 | 0.55 | *HLA-DQB1, HCG9, LINC00266-1* |
| Sweden (exome) | 1,288,701 input variants of 21,390 genes | Schizophrenia | 4969 Cases | 6245 Controls | 0.65 | 0.65 | 0.73 | 0.74 | *ZNF773, PCNT, DYSF* |

*Supplementary Table 1. Summary of the experiments and results in this study for the simplest network in our framework that contains the input SNPs, the gene layer and the output layer. Genes were annotated using ANNOVAR.[2] Manhattan plots for gene importance can be found in Supplementary Materials 2,3 & 4. *MC1R was not present in gene annotations but was identified by linkage disequilibrium. **Many genes contributed to the prediction without clear separation between genes, see Supplementary Materials 4.*

#### 2.2 Pathway networks

| Dataset (type) | Trait | Subjects & phenotype | | AUC GenNet pathway (val) | AUC GenNet pathway (test) | GenNet: top three most important pathways | | |
|---|---|---|---|---|---|---|---|---|
| | | Class I | Class II | | | Global | Mid | Local |
| Rotterdam (genotype array) | Eye color | 4041 Blue | 2250 Other | 0.52 | 0.50 | Organismal Systems (78.4%), Cellular Processes (17.9%), Human Diseases (3.0%) Systems (11.7%) | Digestive system (72.6%), Transport and catabolism (17.9%), Circulatory system (5.7%) | Pancreatic secretion (59.1%), Vitamin digestion and absorption (13.3%), Autophagy - animal (9.5%) |
| UK Biobank (exome) | Hair color | 4501 Blond | 4518 Other | 0.57 | 0.58 | Organismal Systems (70.2%), Environmental Information Processing (21.4%), Cellular Processes (3.5%) | Endocrine system (30.1%), Signal transduction (19.6%), Immune system (14.8%) | Adrenergic signaling in cardiomyocytes (4.1%), Olfactory transduction (3.9%), Insulin signaling pathway (3.6%) |
| | | 15684 Dark brown | 15918 Other | 0.55 | 0.56 | Human Diseases (37.2%), Metabolism (27.2%), Cellular Processes (26.1%) | Metabolism of cofactors and vitamins (26.7%), Substance dependence (17.9%), Transport and catabolism (17.2%) | Thiamine metabolism (26.2%), Endocytosis (17.0%), Parkinson disease (9.6%) |
| | | 1734 Red | 1727 Other | 0.77 | 0.77 | Genetic Information Processing (87.4%), Human Diseases (9.6%), Metabolism (2.8%) | Replication and repair (83.4%), Translation (2.5%), Infectious diseases: Bacterial (2.3%) | Fanconi anemia pathway (79.7%), Homologous recombination (2.2%), Legionellosis (2.0%) |
| | | 16208 Light brown | 16029 Other | 0.57 | 0.57 | Environmental Information Processing (55.1%), Human Diseases (28.3%), Cellular Processes (8.6%) | Signal transduction (54.3%), Infectious diseases: Bacterial (7.8%), Cancers: Overview (5.9%) | MAPK signaling pathway (8.1%), Rap1 signaling pathway (7.2%), Calcium signaling pathway (6.1%) |
| | | 3762 Black | 3753 Other | 0.78 | 0.76 | Organismal Systems (46.9%), Human Diseases (29.7%), Cellular Processes (16.9%) | Endocrine system (18.6%), Cellular community - eukaryotes (12.7%), Nervous system (10.9%) | Axon guidance (5.0%), Focal adhesion (3.7%), Tight junction (3.1%) |
| | Breast cancer | 1070 Cases | 20545 Controls | 0.56 | 0.51 | Human Diseases (57.1%), Organismal Systems (27.1%), Metabolism (8.5%) | Infectious diseases: Viral (16.6%), Cancers: Overview (16.4%), Cancers: Specific types (14.1%) | Pathways in cancer (6.5%), Metabolic pathways (4.2%), Gastric cancer (3.4%) |
| | Diabetes | 2557 Cases | 2555 Controls | 0.54 | 0.54 | Environmental Information Processing (43.5%), Organismal Systems (26.1%), Human Diseases (18.3%) | Signal transduction (40.5%), Nervous system (8.4%), Cancers: Specific types (6.9%) | Ras signaling pathway (7.9%), cAMP signaling pathway (7.2%), MAPK signaling pathway (5.9%) |
| | Atrial fibrillation | 192 Cases | 194 Controls | 0.63 | 0.57 | Organismal Systems (39.6%), Environmental Information Processing (21.0%), Human Diseases (18.6%) | Signal transduction (11.2%), Transport and catabolism (10.7%), Immune system (9.7%) | Cytokine-cytokine receptor interaction (4.4%), Endocytosis (3.8%), Axon guidance (2.9%) |

| | | | | | | | | |
|---|---|---|---|---|---|---|---|---|
| | Coronary Artery Disease | 1563 Cases | 1600 Controls | 0.56 | 0.54 | Environmental Information Processing (29.5%), Organismal Systems (23.7%), Cellular Processes (15.8%) | Signal transduction (27.7%), Cellular community - eukaryotes (8.8%), Immune system (5.5%) | PI3K-Akt signaling pathway (4.6%), Focal adhesion (3.7%), Tight junction (2.5%) |
| | Bipolar disorder | 343 Cases | 347 Controls | 0.55 | 0.47 | Organismal Systems (76.9%), Cellular Processes (16.4%), Metabolism (5.7%) | Endocrine system (46.5%), Transport and catabolism (16.4%), Immune system (11.8%) | Melanogenesis (32.9%), Thyroid hormone signaling pathway (12.2%), Phagosome (9.0%) |
| | Dementia | 139 Cases | 142 Controls | 0.58 | 0.55 | Human Diseases (39.8%), Environmental Information Processing (26.5%), Organismal Systems (22.8%) | Signal transduction (22.2%), Cancers: Overview (13.3%), Endocrine system (7.3%) | Pathways in cancer (5.6%), PI3K-Akt signaling pathway (4.1%), Proteoglycans in cancer (2.8%) |
| | Male balding pattern | 3454 Balding pattern 1 | 3454 Balding pattern 4 | 0.55 | 0.54 | Organismal Systems (34.6%), Human Diseases (22.7%), Metabolism (20.8%) | Nervous system (9.7%), Global and overview maps (8.9%), Transport and catabolism (8.3%) | Metabolic pathways (8.7%), Endocytosis (4.3%), Axon guidance (4.1%) |
| | Asthma | 4229 Cases | 4214 Controls | 0.54 | 0.51 | Genetic Information Processing (52.3%), Metabolism (23.6%), Human Diseases (23.2%) | Folding, sorting and degradation (41.5%), Cardiovascular diseases (12.6%), Amino acid metabolism (12.2%) | Ubiquitin mediated proteolysis (22.0%), Protein processing in endoplasmic reticulum (11.8%), RNA transport (10.8%) |
| Sweden (exome) | Schizophrenia | 4969 Cases | 6245 Controls | 0.67 | 0.68 | Human Diseases (30.8%), Organismal Systems (26.7%), Genetic Information Processing (26.5%) | Infectious diseases: Viral (27.3%), Endocrine system (16.6%), Folding, sorting and degradation (13.7%) | Human papillomavirus infection (11.7%), Ubiquitin mediated proteolysis (10.0%), Ribosome biogenesis in eukaryotes (4.4%) |

*Supplementary Table 2: Neural networks build with gene and KEGG pathway[3] information. The pathway information is hierarchical with 3 levels: global, mid and local. In this table the top three pathways for each level are displayed in terms of relative contribution. Interactive plots for all phenotypes can be found at: https://github.com/ArnovanHilten/GenNet_paper_plots*

#### 2.2 GTEx expression networks

| Dataset (type) | Number of input variants | Trait | Subjects & phenotype | | AUC GenNet GTEx (val) | AUC GenNet GTEx (test) | GenNet: top three most important cell types |
|---|---|---|---|---|---|---|---|
| | | | Class I | Class II | | | |
| Rotterdam (genotype array) | 113,241 (exonic) inputs of 16,628 genes | Eye color | 4041 Blue | 2250 Other | 0.76 | 0.76 | Esophagus Gastroesophageal Junction (10.1%), Colon Sigmoid (9.5%), Esophagus Muscularis (7.5%) |
| UK Biobank (exome) | 6,986,636 input variants of 15,827 genes | Hair color | 4501 Blond | 4518 Other | 0.67 | 0.64 | Brain Nucleus accumbens (basal ganglia) (4.1%), Brain Putamen (basal ganglia) (3.6%), Artery Tibial (3.5%) |
| | | | 15684 Dark brown | 15918 Other | 0.60 | 0.62 | Brain Frontal Cortex (BA9) (4.3%), Brain Hippocampus (3.6%), Brain Caudate (basal ganglia) (3.5%) |
| | | | 1734 Red | 1727 Other | 0.76 | 0.76 | Brain Caudate (basal ganglia) (4.1%), Esophagus Muscularis (4.1%), Brain Amygdala (3.9%) |
| | | | 16208 Light brown | 16029 Other | 0.61 | 0.61 | Brain Putamen (basal ganglia) (3.9%), Brain Hypothalamus (3.7%), Brain Cerebellar Hemisphere (3.6%) |
| | | | 3762 Black | 3753 Other | 0.80 | 0.80 | Artery tibial (4,0%), artery aorta (3.7%), esophagus gastroesophageal junction (3.7%), |
| | | Diabetes | 2557 Cases | 2555 Controls | 0.55 | 0.56 | Brain Cortex (4.4%), Esophagus Gastroesophageal Junction (4.4%), Artery Tibial (4.2%) |
| | | Atrial fibrillation | 192 Cases | 194 Controls | 0.65 | 0.49 | Esophagus Muscularis (4.2%), Brain Cerebellum (4.1%), Artery Aorta (3.6%) |
| | | Coronary Artery Disease | 1563 Cases | 1600 Controls | 0.55 | 0.54 | Brain Caudate (basal ganglia) (4.2%), Brain Putamen (basal ganglia) (4.1%), Brain Amygdala (4.0%) |
| | | Bipolar disorder | 343 Cases | 347 Controls | 0.56 | 0.58 | Brain Caudate (basal ganglia) (4.6%), Brain Hippocampus (4.6%), Brain Cerebellar Hemisphere (4.3%) |
| | | Dementia | 139 Cases | 142 Controls | 0.52 | 0.41 | Brain Putamen (basal ganglia) (4.3%), Brain Caudate (basal ganglia) (4.2%), Brain Amygdala (4.1%) |
| Sweden (exome) | 1,288,701 input variants of 21,390 genes | Schizophrenia | 4969 Cases | 6245 Controls | 0.66 | 0.66 | Uterus (4.2%), Colon Sigmoid (3.8%), Breast Mammary Tissue (3.7%) |

*Supplementary Table 3: Neural networks build with genes and tissue expression (GTEx) only the significantly different expressed genes are connected to their tissues Finucane et al. (2018)[4] to ensure an interpretable network. Interactive plots for all phenotypes can be found at:*

*https://github.com/ArnovanHilten/GenNet_paper_plots*

#### 2.2 ImmGen expression networks

| **Dataset (type)** | **Trait** | **Subjects & phenotype** | | **AUC GenNet GTEx (val)** | **AUC GenNet GTEx (test)** | **Most important immunological cell types** |
|---|---|---|---|---|---|---|
| | | **Class I** | **Class II** | | | |
| Rotterdam (genotype array) | Eye color | 4041 Blue | 2250 Other | 0.50 | 0.50 | Tgd.vg2+24ahi.Th (5.3%), T.DPsm.Th (5.0%), Ep.5wk.MEChi.Th (4.1%) |
| UK Biobank (exome) | Hair color | 4501 Blond | 4518 Other | 0.50 | 0.48 | proB.FrA.BM (5.5%), MF.Microglia.CNS (4.9%), SC.CDP.BM (4.7%) |
| | | 15684 Dark brown | 15918 Other | 0.50 | 0.50 | T.8SP24int.Th (0.6%), MF.F480hi.ctrl.PC (0.6%), DN.SLN.CFA.d6.v2 (0.6%) |
| | | 1734 Red | 1727 Other | 0.49 | 0.47 | DC.8+.MLN (2.1%), GN.BM (2.1%), DC.8-4-11b-.MLN (1.8%) |
| | | 16208 Light brown | 16029 Other | 0.48 | 0.50 | FRC.SLN (0.6%), Tgd.vg5-.act.IEL (0.6%), MEChi.GFP+.Adult.KO (0.6%) |
| | | 3762 Black | 3753 Other | 0.79 | 0.78 | T.8SP24int.Th (0.8%), T.4Mem44h62l.LN (0.8%), T.4Mem.LN (0.8%) |
| | Male balding pattern | 3454 Balding pattern 1 | 3454 Balding pattern 4 | 0.50 | 0.51 | GN.Bl.v2 (1.4%), DC.103+11b-.Lv (1.4%), DC.8-4-11b+.Sp (1.3%) |
| | Atrial fibrillation | 192 Cases | 194 Controls | 0.53 | 0.41 | Ep.5wk.MEClo.Th (0.8%), T.4.Pa.BDC (0.8%), Ep.MEChi.Th (0.8%) |
| | Coronary Artery Disease | 1563 Cases | 1600 Controls | 0.57 | 0.53 | Tgd.vg2+24ahi.Th (0.8%), DC.103-11b+F4/80lo.Kd (0.8%), Tgd.vg2+.Sp (0.8%) |
| | Bipolar disorder | 343 Cases | 347 Controls | 0.54 | 0.57 | FRC.Cad11.WT.v2 (0.8%), T.4Mem.Sp (0.8%), CD19Control (0.8%) |
| | Dementia | 139 Cases | 142 Controls | 0.54 | 0.57 | FRC.Cad11.WT.v2 (0.8%), DN.SLN.v2 (0.8%), preB.FrD.BM (0.8%) |
| Sweden (exome) | Schizophrenia | 4969 Cases | 6245 Controls | 0.73 | 0.74 | FRC.MLN (1.2%), DN.SLN.v2 (1.1%), SC.LT34F.BM (1.1%) |

*Supplementary Table 4: Neural networks build with genes and Immunological Genome Project data*[5]*. Only the significantly different expressed genes are connected to their tissues Finucane et al. (2018)*[4] *to ensure an interpretable network. Interactive plots for all phenotypes can be found at: https://github.com/ArnovanHilten/GenNet_paper_plots*

**Supplementary 3. Sweden-Schizophrenia population-based case-control study**

All networks applied on the Sweden Schizophrenia Case Control study are trained with similar parameters, found by optimizing the performance on the validation set for the 'gene network'. The batch size was chosen to be 200, the optimizer Adam (learning rate of 0.0006), and with batch normalization after every tangent hyperbolicus activation. All networks are trained on a single GPU (Nvidia Geforce 1080 GTX) and converged within 3 hours.

| Network | AUC val. set | AUC test set | Accuracy test set | Source |
|---|---|---|---|---|
| Gene Network | 0.73 | 0.74 | 0.68 | Annovar gene annotations |
| Gene Network L1 | 0.75 | 0.74 | 0.68 | Annovar gene annotations |
| Gene -Pathway | 0.67 | 0.68 | 0.61 | Kegg pathway annotations |
| Gene-Tissue expr. | 0.66 | 0.66 | 0.60 | GTEx tissue expr. |
| Gene-Cell. expr. | 0.74 | 0.75 | 0.68 | FUMA |

*Supplementary Table 5. Overview of the performance for different networks. Area Under the Curve (AUC), Precision, Accuracy and F1 score for the different types of networks. The theoretical maximum accuracy is ~ 72%. The dataset was split 60/20/20 in training validation and with respect to the ratio of cases and controls. All results are reported on the test set with the optimum decision threshold determined by the ROC curve of the validation set.*

#### 3.1 Gene network

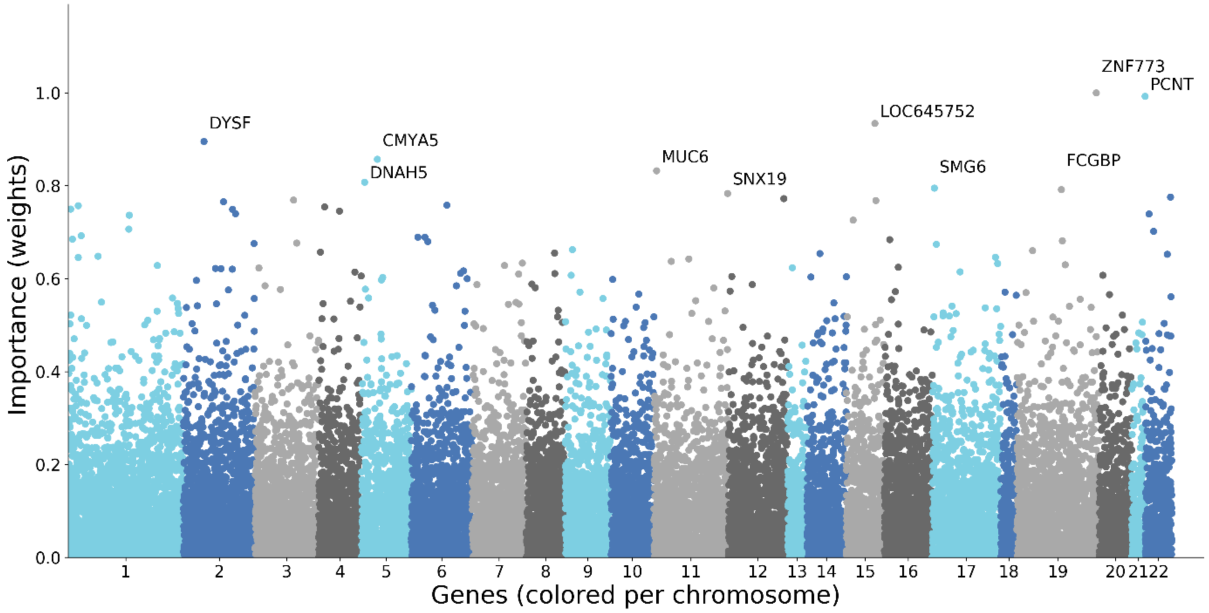

Supplementary Figure 4. Gene Importance for Schizophrenia predictions. Area under the curve for this model was 0.73 in the validation set and 0.74 in the test set. This was the best run of a series of experiments run with the same parameters. These models obtained a mean AUC of 0.70 ± 0.018 in the validation set and 0.72 ± 0.016 in the test set.

##### 3.2 Gene network with an L1 constraint on the weights

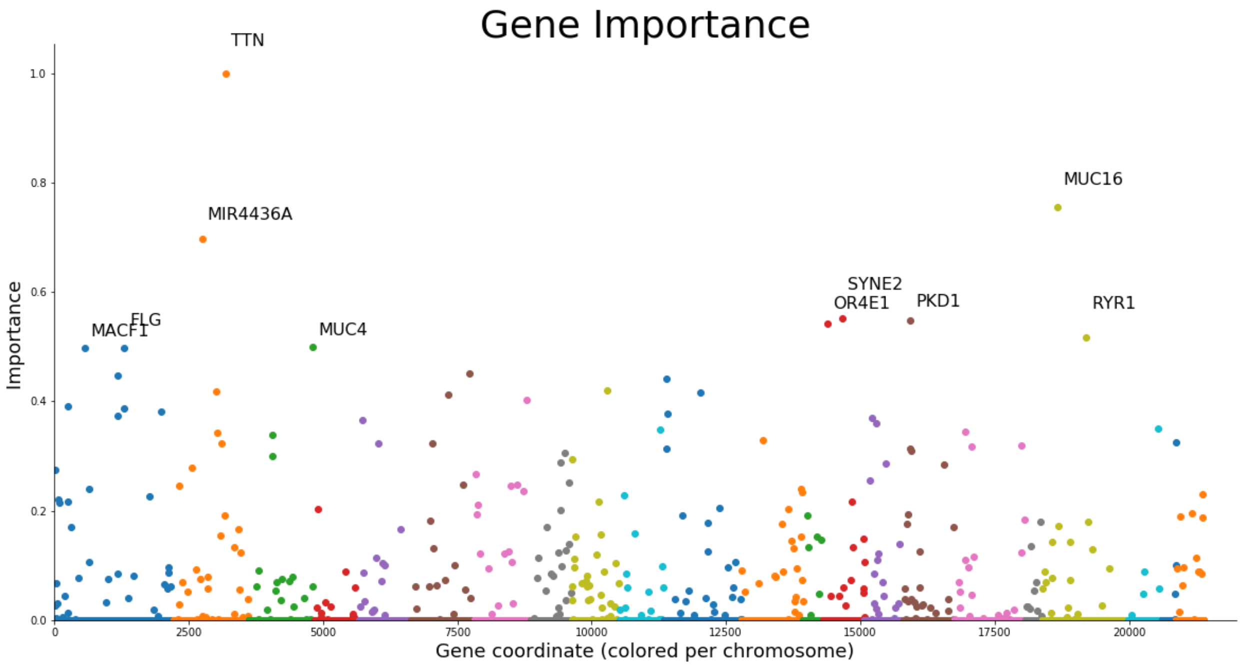

Supplementary Figure 5. Similar to lasso logistic regression, the network can be forced to focus on only the most important genes by adding L1 regularization, easing interpretation while maintaining performance (AUC of 0.74). The network is forced to focus only on the most important genes, leaving most genes with a near zero weight. As a consequence, larger genes are selected, such as *TTN* and *MUC16*. *MIR4436A* is therefore interesting since this is a micro-RNA of 63 base pairs.

##### 3.3 Pathway network

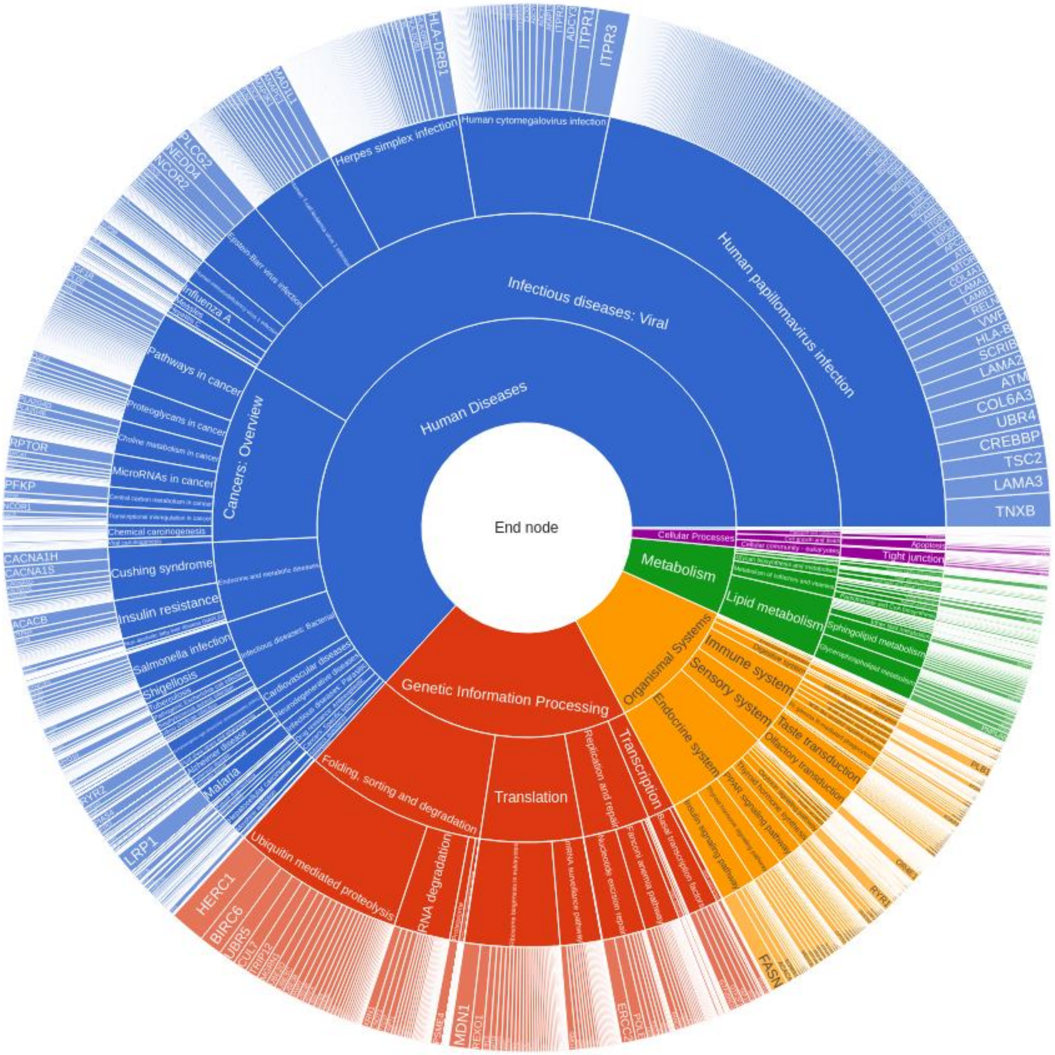

Supplementary Figure 6. The gene-pathway network achieved an AUC of 0.68 in the test set with only 7204 genes annotated by the KEGG database, leaving more than 14 000 genes unconnected. The performance of this network could thus be artificially inflated due to the same LD structure, leaking information of unconnected genes without a pathway. Pathway nodes are not as objectively and uniquely defined as gene nodes (see section 2.4). Nonetheless, pathway annotations could possibly regulate the network further to assign high weights to important pathways, genes and SNPs, guiding the network to select only disease relevant entities. This would ease interpretability, avoid tagging and will definitely increase performance. Moreover, pathway annotations allow us to create deeper networks, carrying more information about how;

from important SNPs to genes, genes to pathways and from pathways to the eventual trait everything relates to each other with respect to the trait.

##### 3.4 Gene-cell network & gene-GTEx networks

There are multiple variations of these networks depending on the threshold used for defining connections (i.e., what is the level of expression required to start defining connections?).
An interesting point of research and a concern for the interpretability is the uniqueness of the connections required to be still interpretable. In the tissue and cell expression the connections are not as unique as in the gene layer, where overlap between inputs for neurons is rare. To avoid an arbitrary threshold the weights of the connections could be initialized with the level of expression. However, there is no guarantee that such a trained network would be interpretable.
To create more uniquely defined nodes, we chose to group and connect only the significantly associated genes per tissue or cell type. For this we used the approach and resources found in Finucane et al (2018)[4]. However, interpretability is still not guaranteed and further research is necessary to confirm interpretability.
Tutorials and examples for creating networks with cell and tissue expression are available on GitHub.

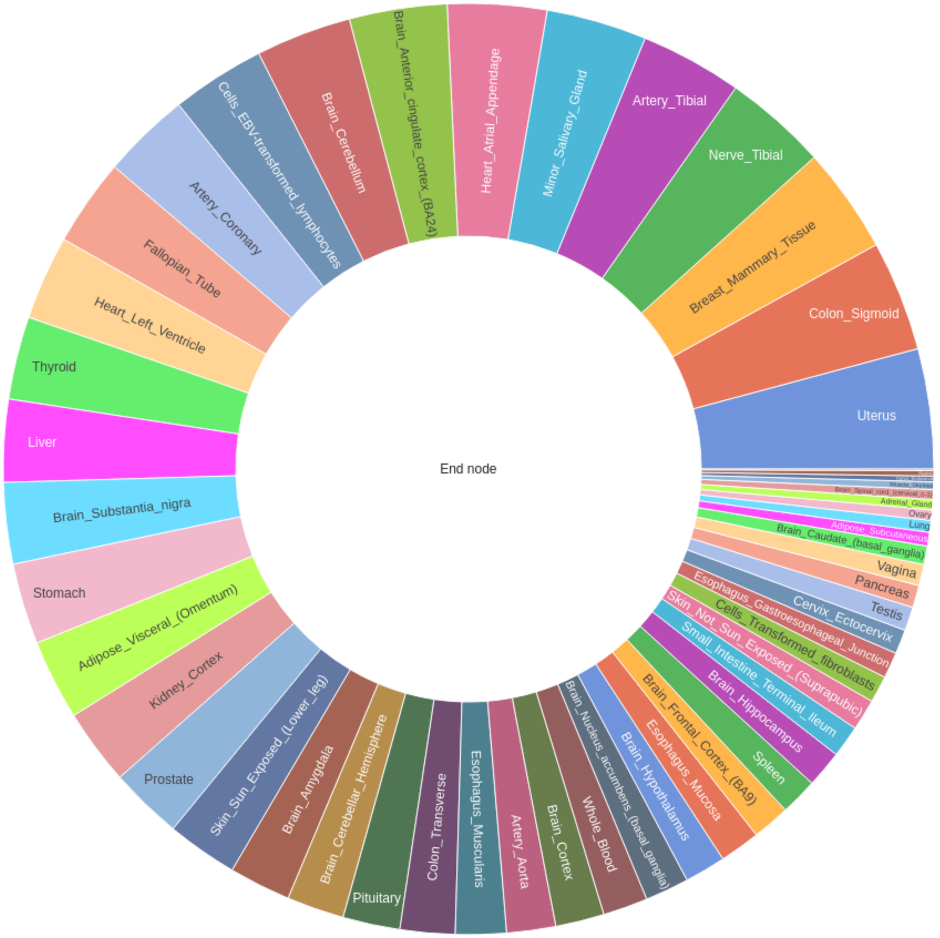

*Supplementary figure 7: Sunburst of the tissues and their relative importance using the GTEx tissue expression layer.*

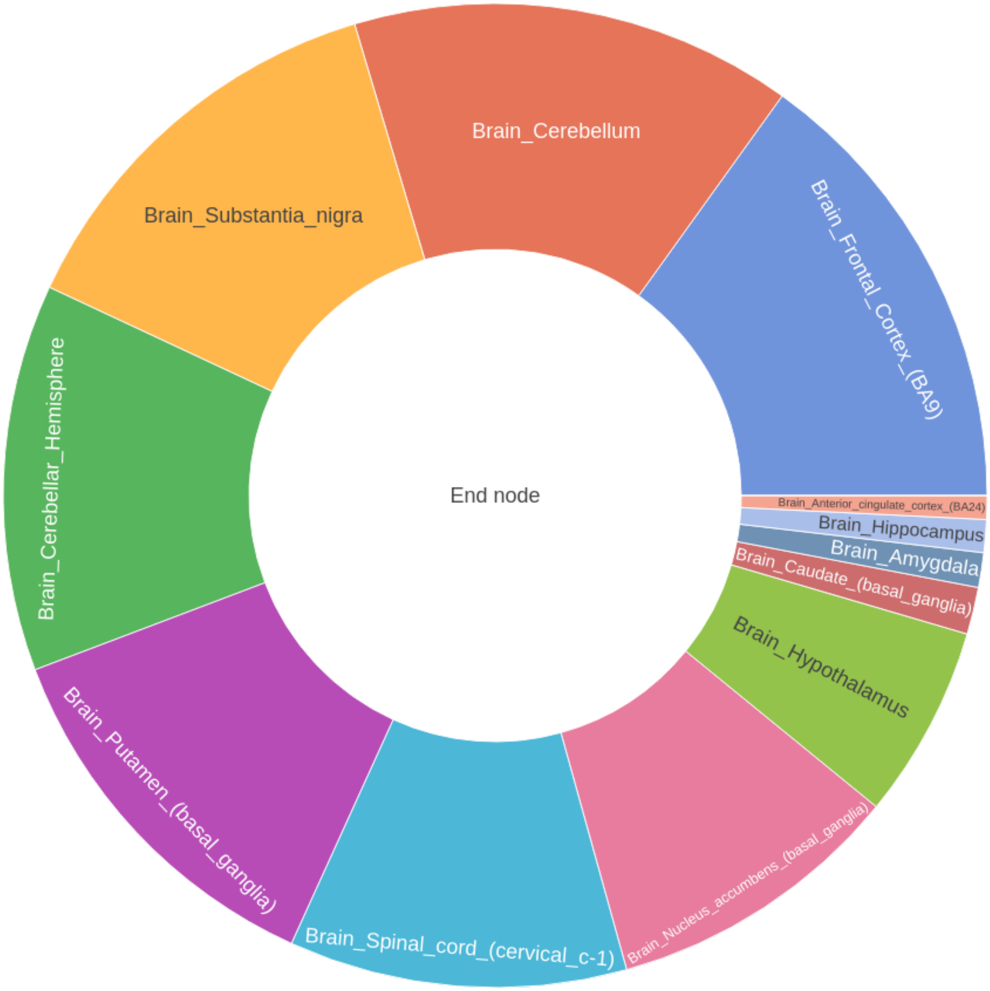

*Supplementary figure 8: Sunburst of the brain tissues and their relative importance using the GTEx brain expression layer.*

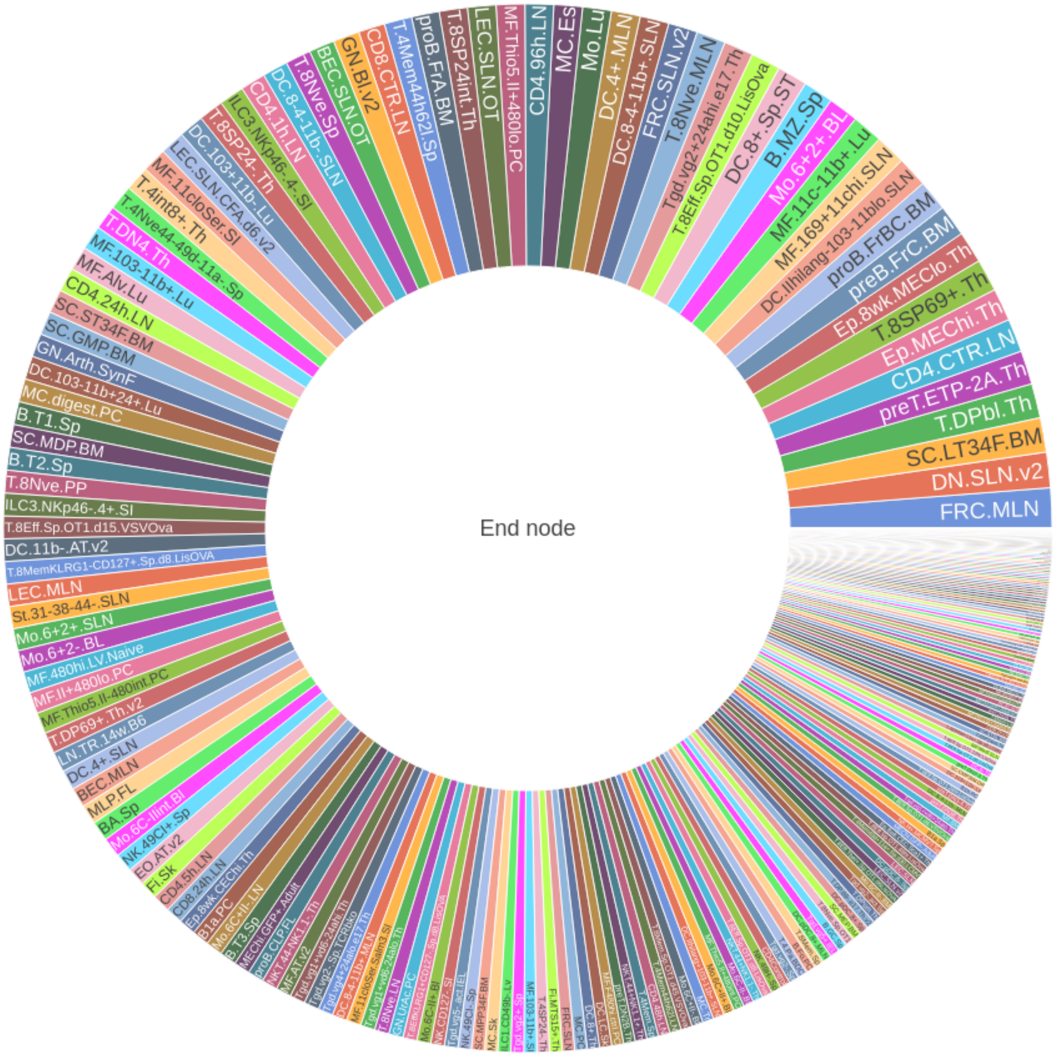

*Supplementary figure 9: Sunburst of the immune cell types and their relative importance using the* ImmGen *layer. We used the preprocessed t-statistics made available from Finucane et al. (2018)* [4] *Raw data can be obtained from (*[http://www.immgen.org/](http://www.immgen.org/) *)*[5]

#### Supplementary 4. UK Biobank

##### 4.1 Bipolar disorder

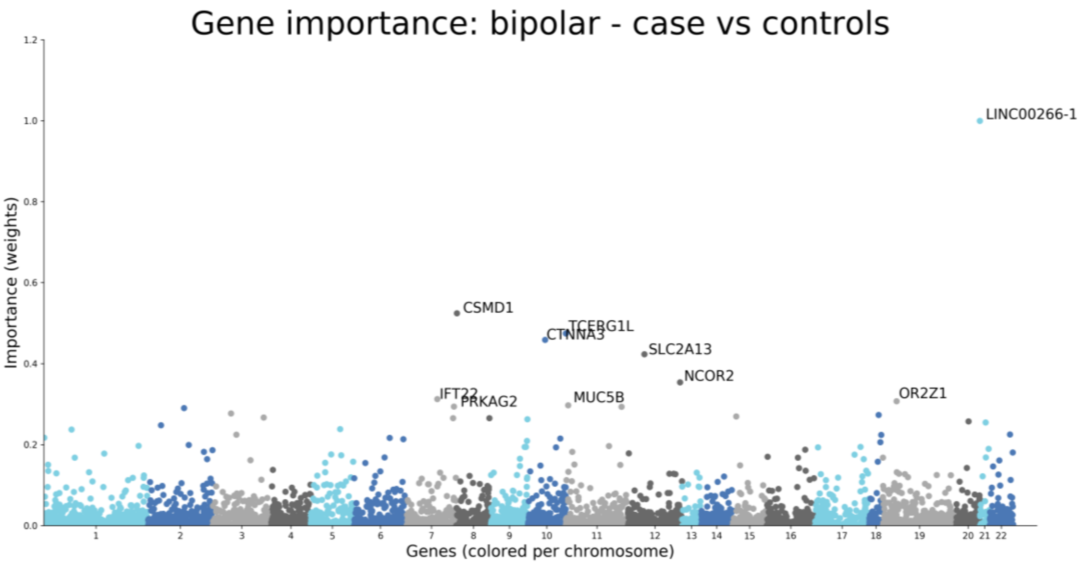

Supplementary Figure 10. Gene contribution for the prediction of bipolar disorder. *LINC00266-1* (Long Intergenic Non-Protein Coding RNA 266-1) on chromosome 22 is the greatest contributor.

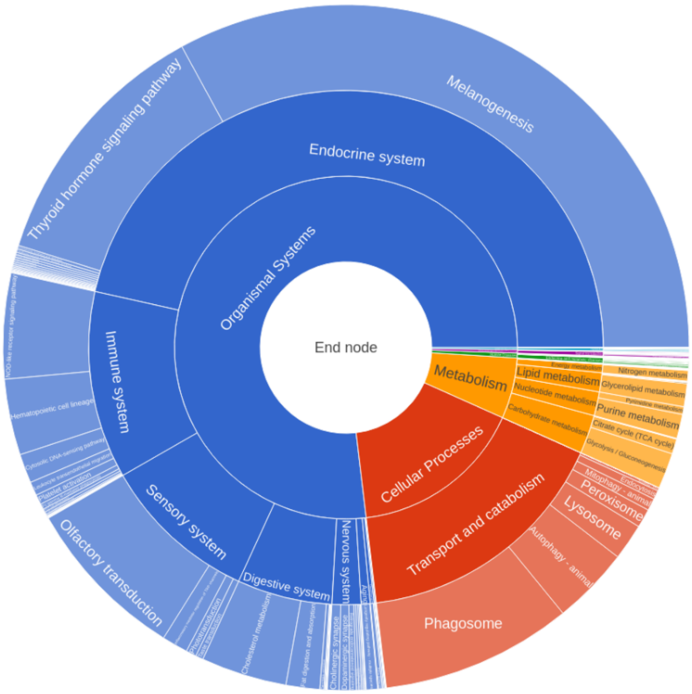

Supplementary Figure 11. Pathway contribution for the prediction of bipolar disorder.

#### 4.2 Skin color

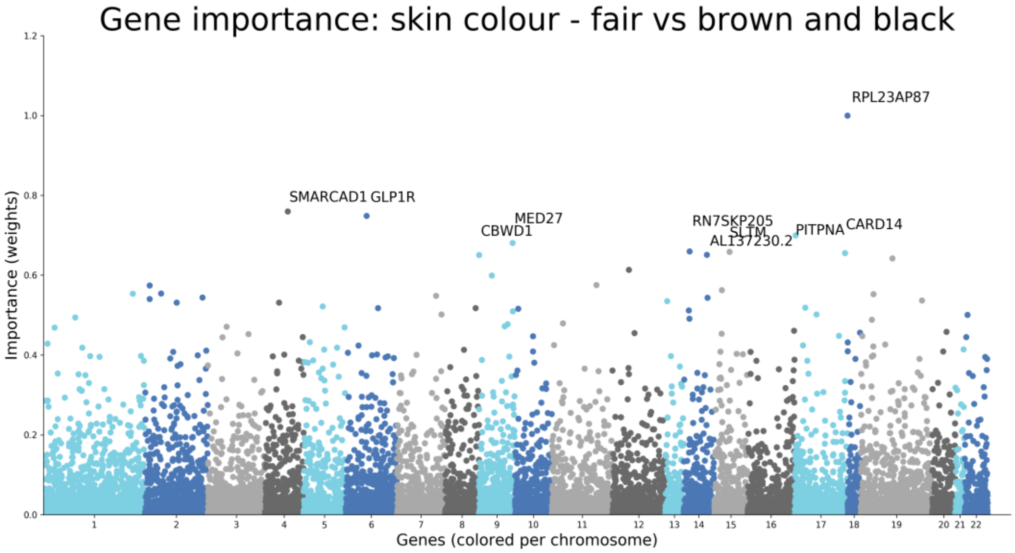

Supplementary Figure 12. The network to predict skin color obtains a near perfect score (AUC of 0.98 in the validation and test set) for distinguishing between a fair or a dark skin color Predicting skin color (fair vs brown and black) is close to predicting ethnicity. The predictions for skin color can serve as a good example why interpretability is a necessity for reliable predictions: interpretation of the network showed that the predictions are based on other factors such as ethnic background rather than genes related to pigmentation and skin color. Neural networks have better predictive capabilities than regular linear methods and can therefore identify and exploit biases in the dataset more easily. While using neural networks we should thus pay even more attention to confounders. *SMARCAD1* is associated with not having fingerprints.[6]

##### 4.3 Hair color: one versus all

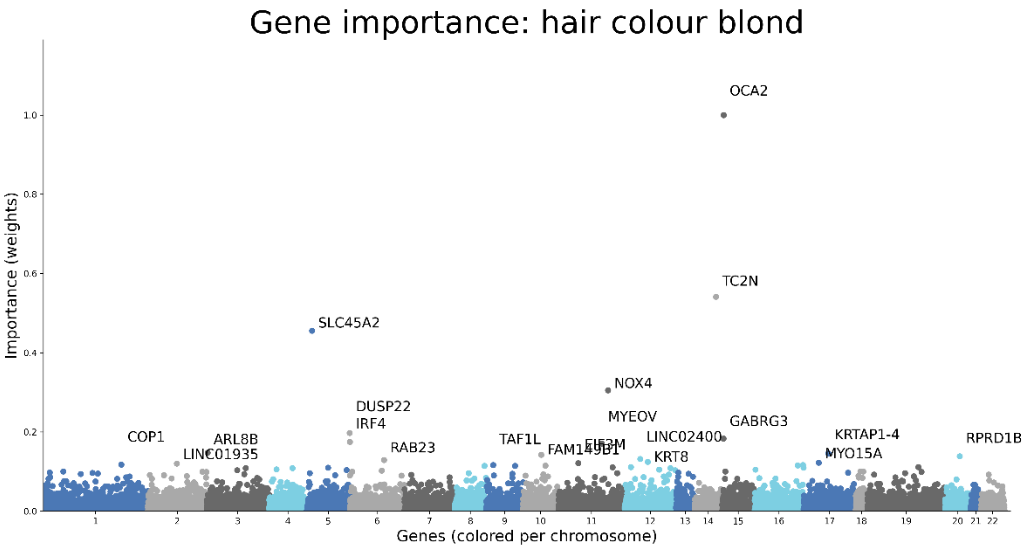

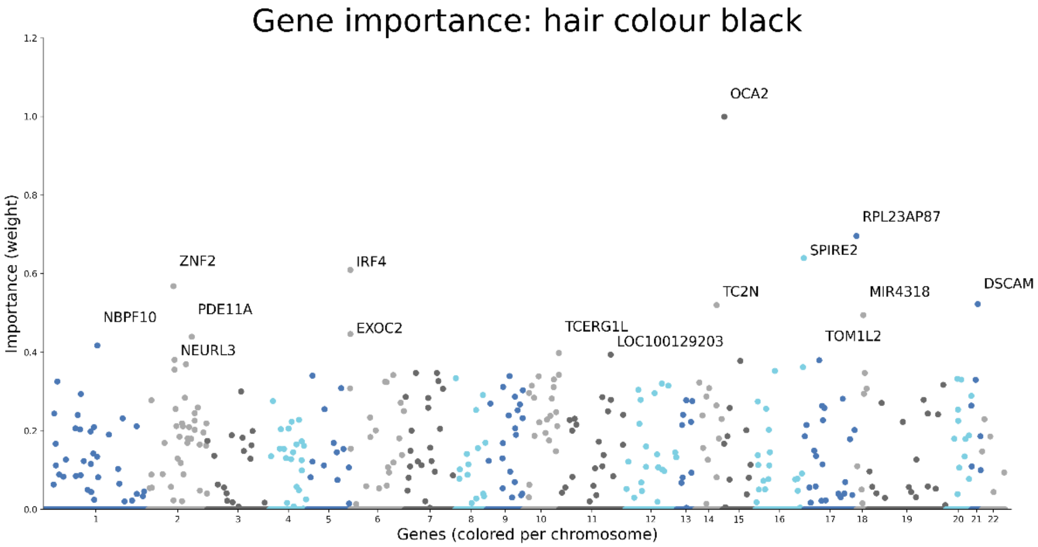

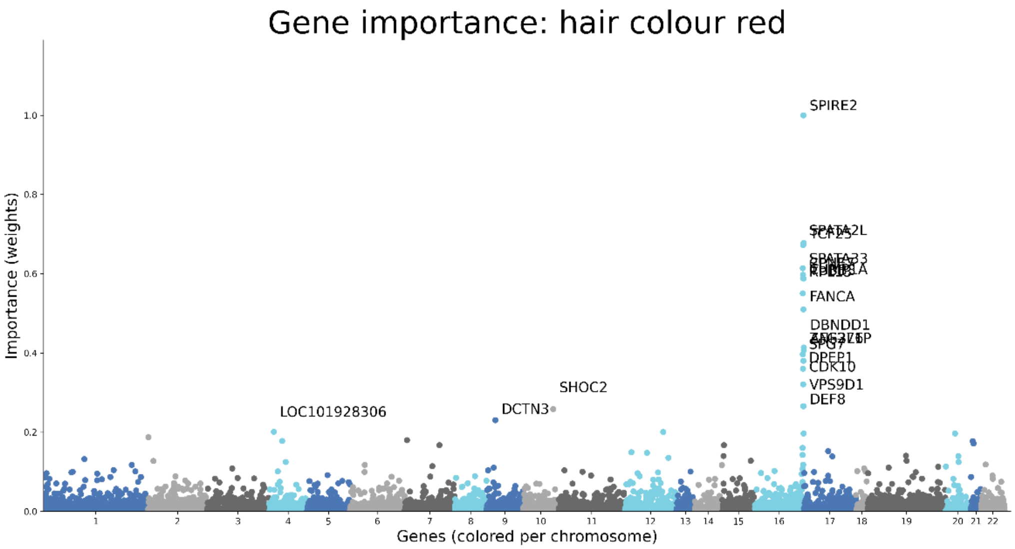

Gene importance: hair colour red
Importance (weights)
Genes (colored per chromosome)
SPIRE2
SPATA2L
SPATA33
FANCA
DBNDD1
SPG7
DPEP1
CDK10
VPS9D1
DEF8
SHOC2
DCTN3
LOC101928306

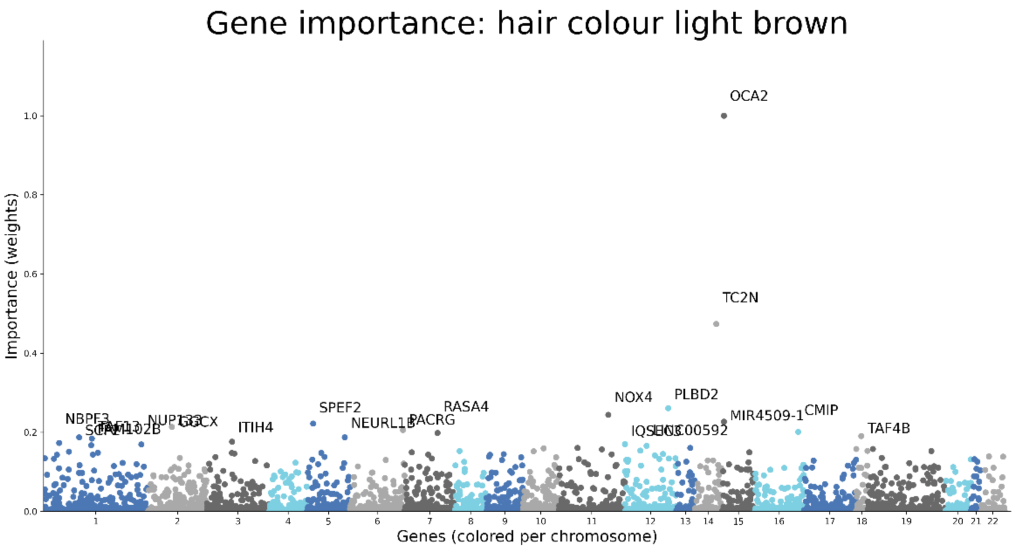

Gene importance: hair colour light brown
Importance (weights)
Genes (colored per chromosome)
OCA2
TC2N
NOX4
PLBD2
MIR4509-1
CMIP
TAF4B
SPEF2
RASA4
NEURL1B
PACRG
ITIH4
NBPF3

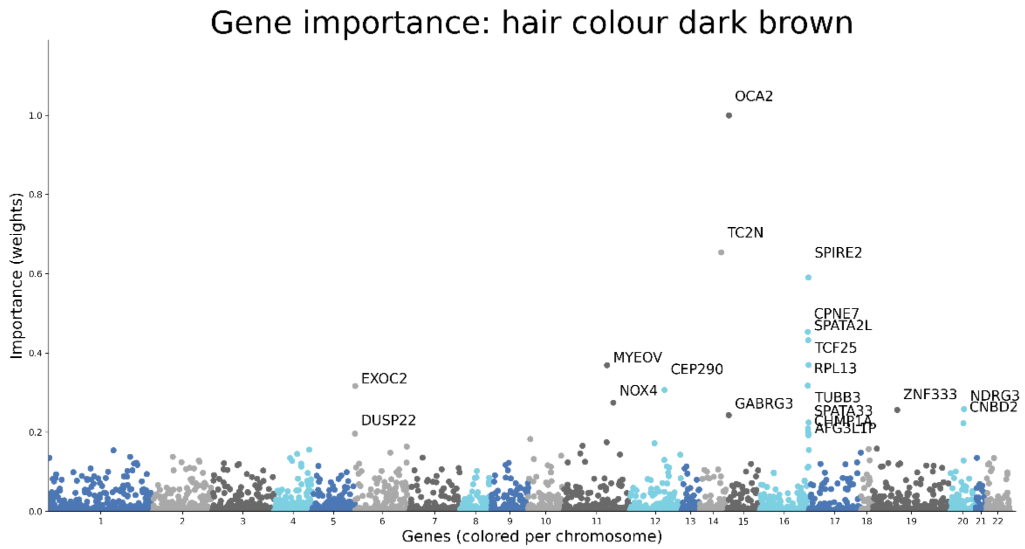

Supplementary Figure 13 to 17: The region on chromosome 16 contains the *MC1R* gene, a well-known gene that is associated with red hair color. Even though this gene was not present in the annotations, LD structure and the interpretability of the network allowed us to identify this gene. *OCA2* is a known gene, a melanin precursor, *EXOC2* has been identified before by Han et al. (2008).[7]

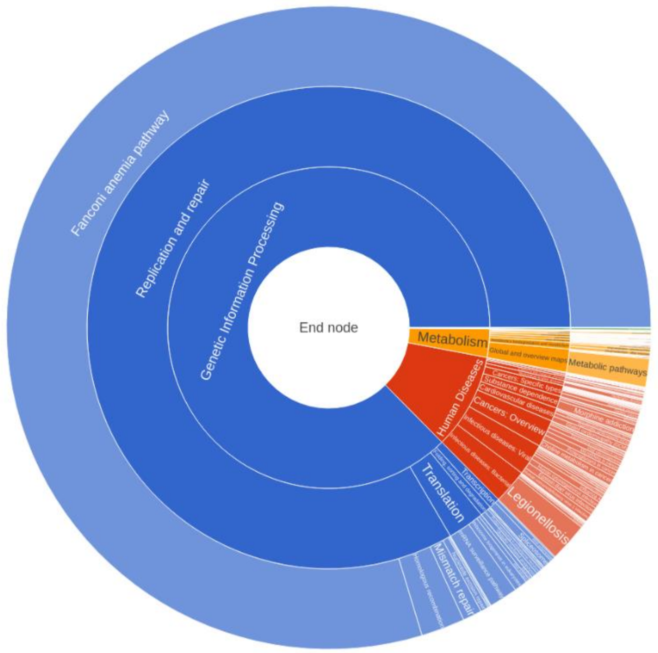

Supplementary Figure 18: Red hair color was dominated by the Fanconi anemia pathway.

#### 4.4 Hair color: one vs one

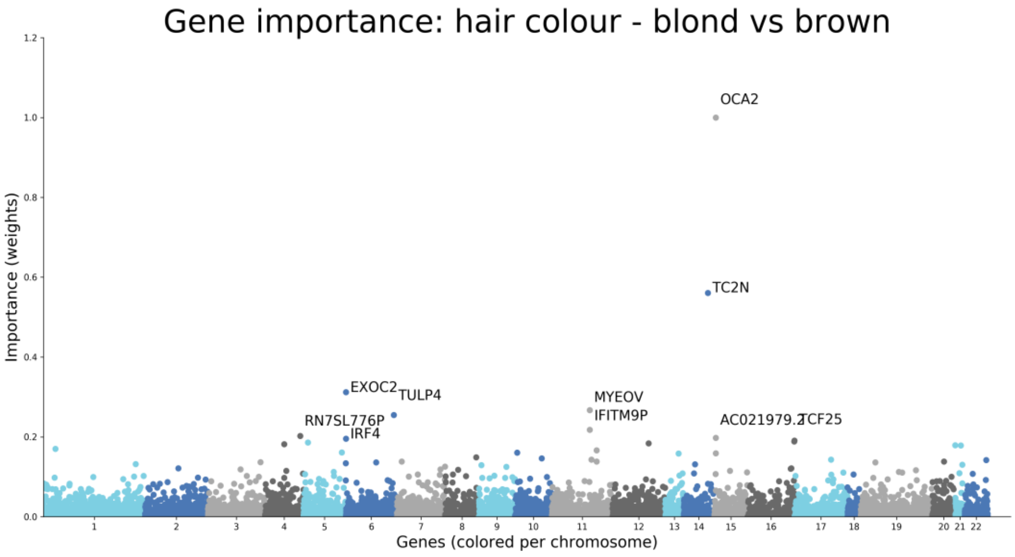

Supplementary Figure 18. *OCA2* is a known gene, a melanin precursor, *EXOC2* has been identified before by Han et al. (2008).[7]

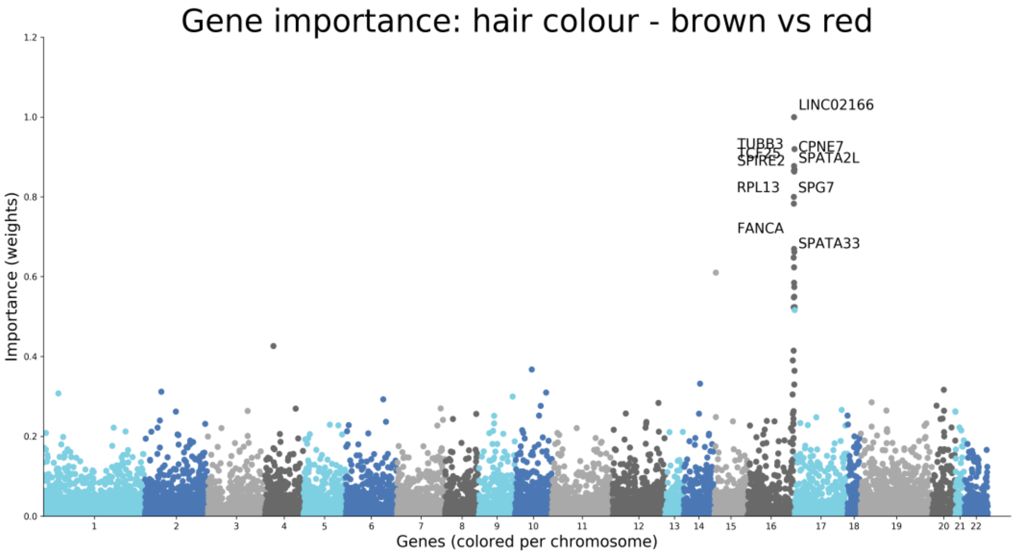

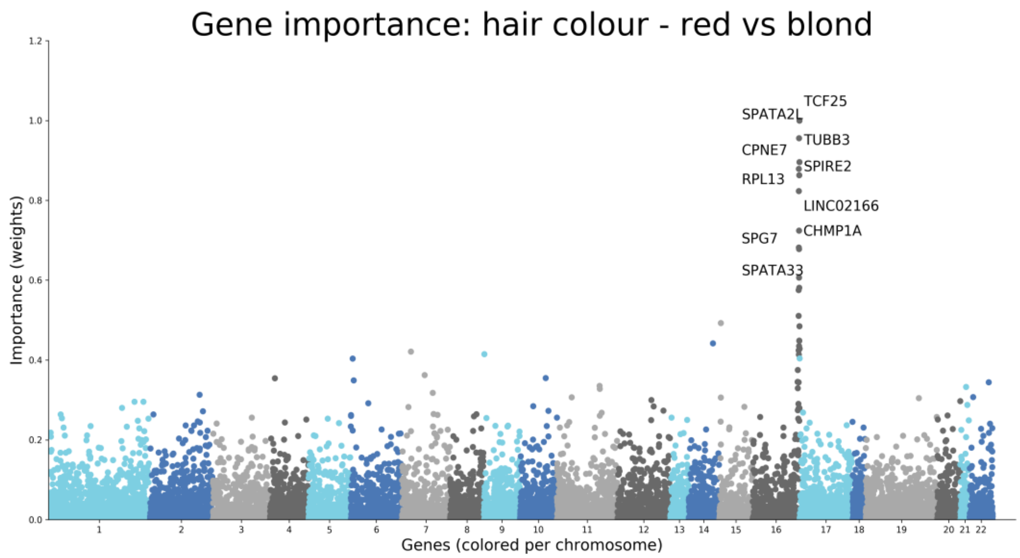

Supplementary Figure 19 & 20: Brown versus red and red versus blond hair. This region on chromosome 16 contains the *MC1R* gene, a well-known gene that is associated with red hair color. Even though this gene was not present in the annotations, LD structure and the interpretability of the network allowed us to identify this gene.

##### 4.5 Male balding pattern

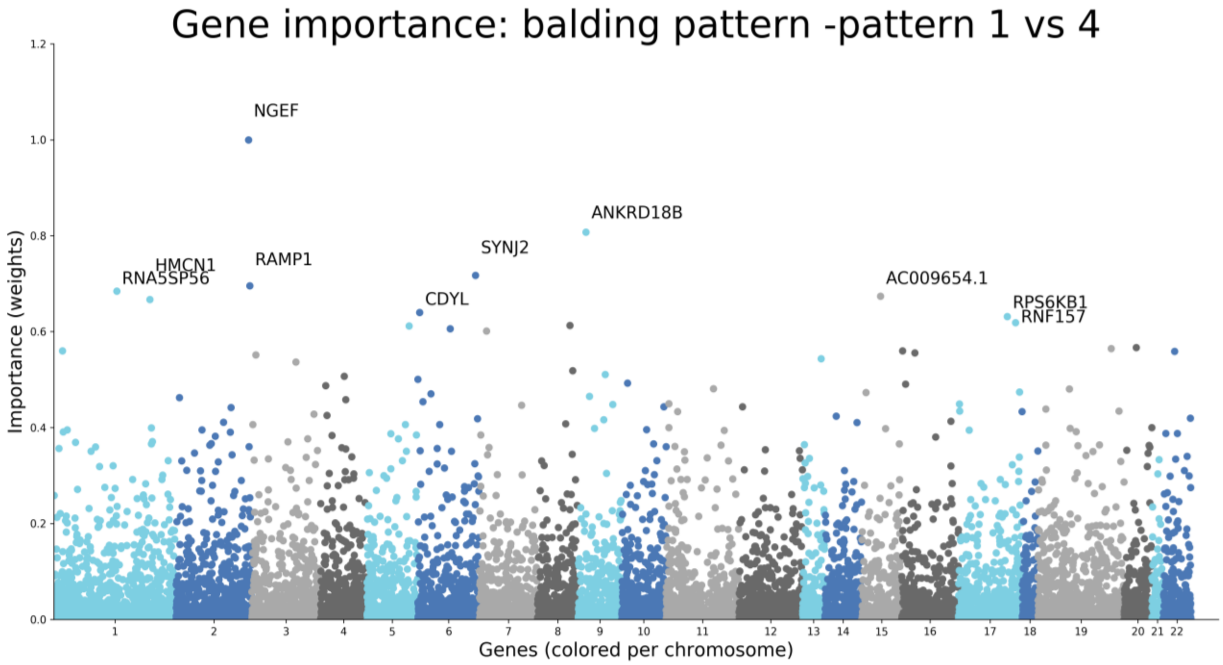

Supplementary Figure 21. Manhattan plot for genes important for the network for separating between balding pattern 1 and 4.

##### 4.6 Breast cancer

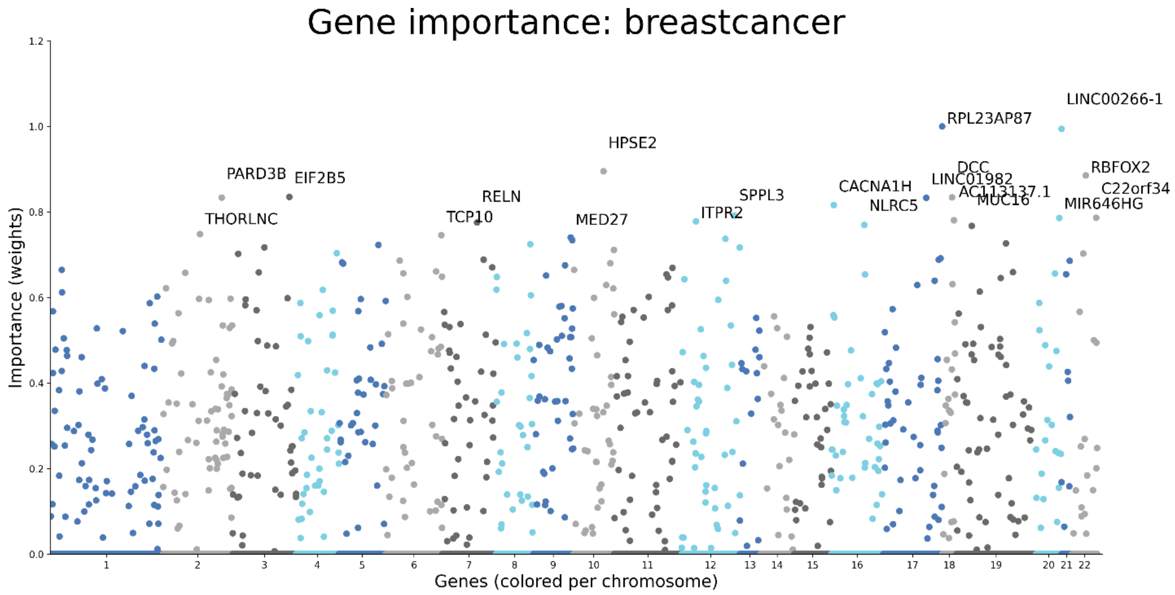

Supplementary Figure 22. Manhattan plot for genes important for the network for separating between cases and controls.

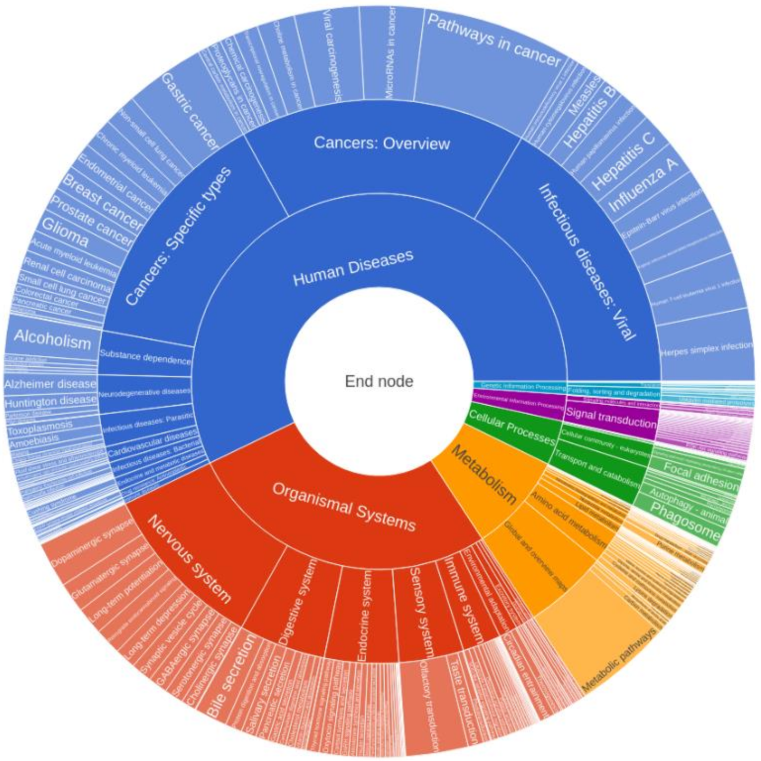

Supplementary Figure 23. Sunburst plot of pathways for cancers. Predictive performance for the pathway network was very low with 0.51 in the test set, 0.56 in the validation set.

##### 4.7 Asthma

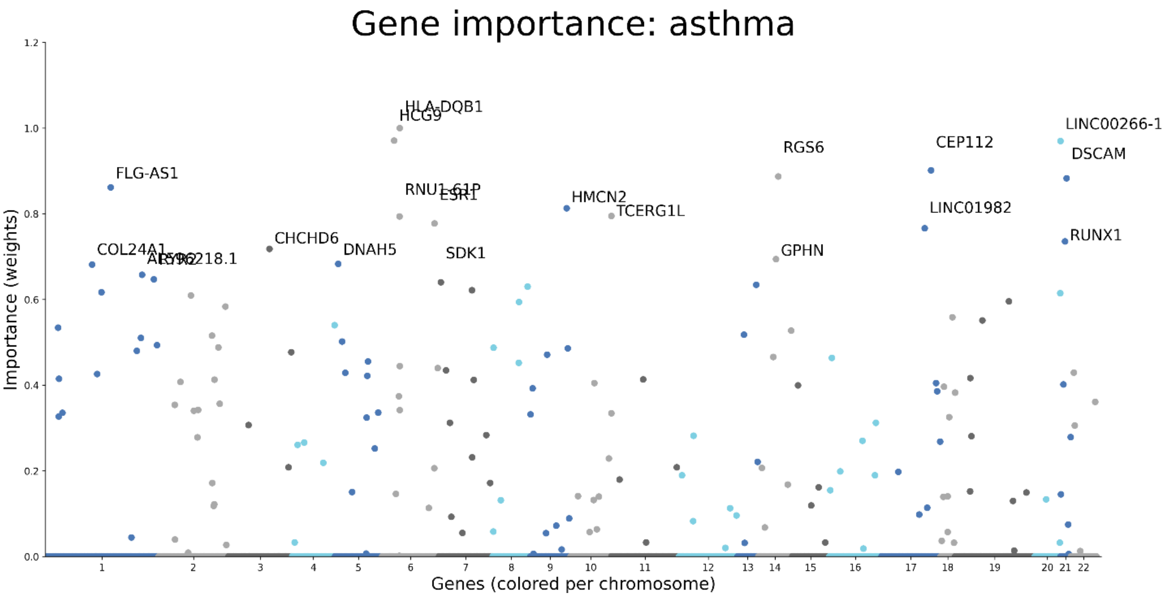

Supplementary Figure 24. Manhattan plot for genes important for the network for separating between cases and controls for asthma.

##### 4.8 Dementia

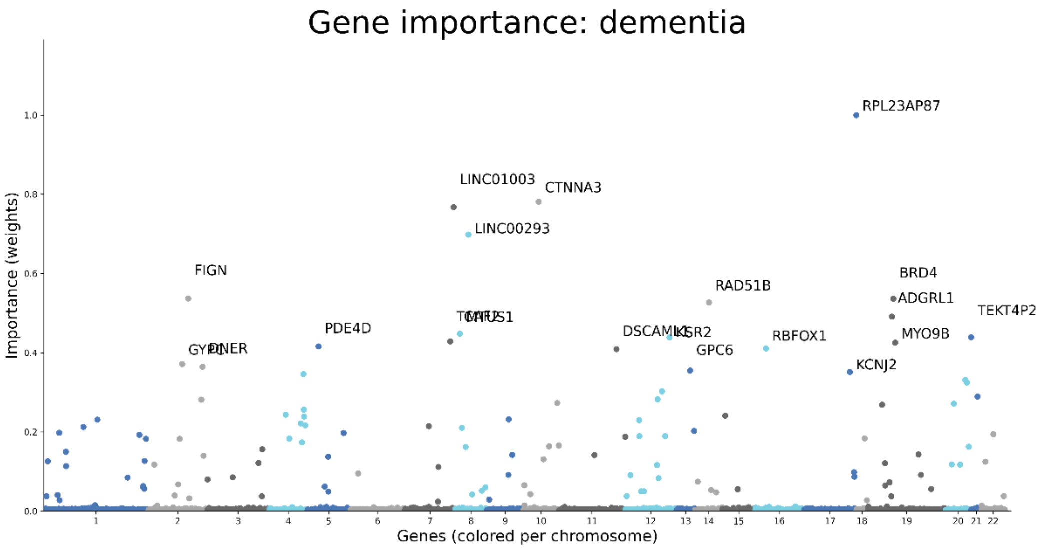

Supplementary Figure 25. Manhattan plot for genes important for the network for separating between cases and controls for dementia. APOE got assigned a (normalized) weight of 0.08

##### 4.9 Coronary artery disease

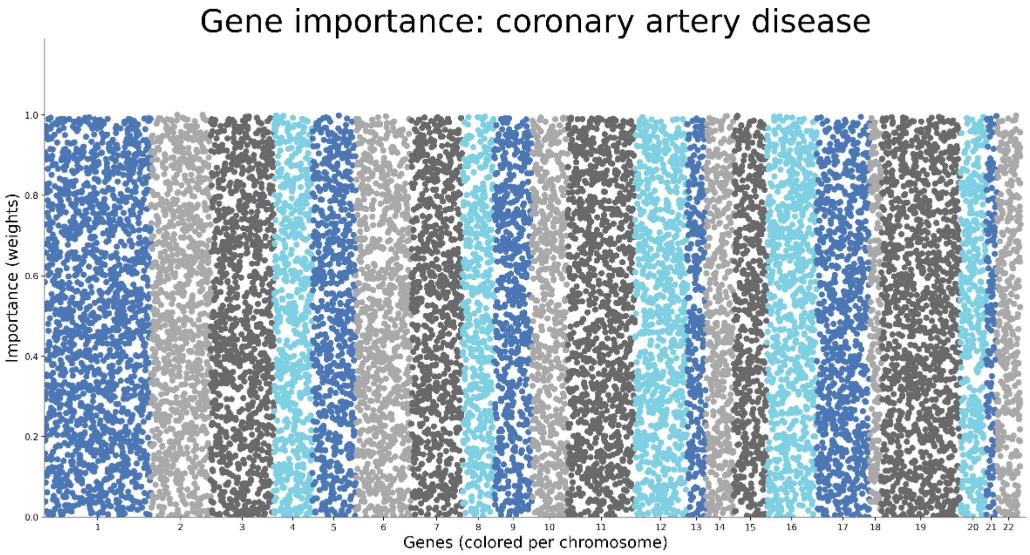

Supplementary Figure 26. Manhattan plot for genes important for the network for separating between cases and controls for coronary artery disease. The predictive performance was poor; 0.56 in the test set and 0.58 in the validation set and we suspect that we are underpowered. More data will most likely improve the predictive performance and therefore the interpretation.

##### 4.10 Atrial fibrillation

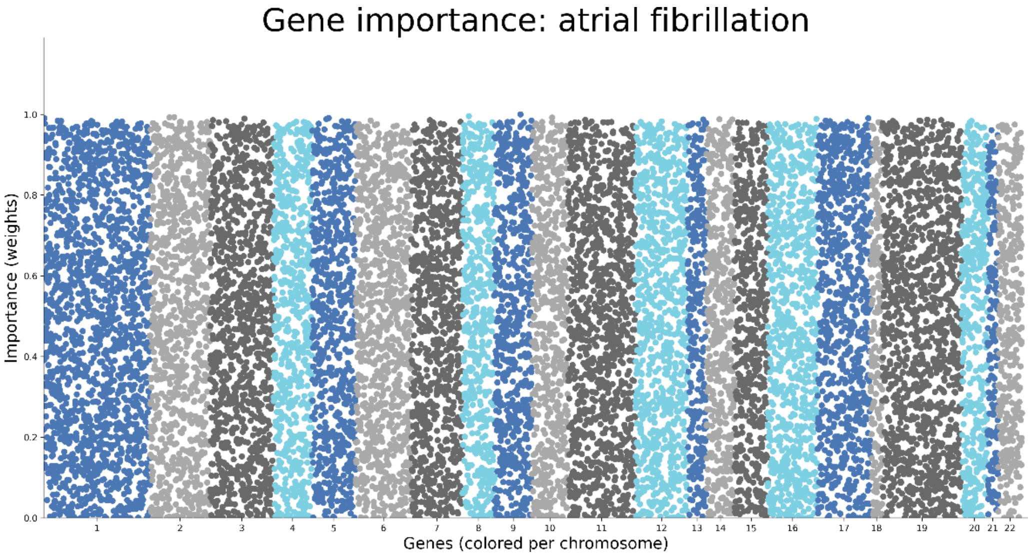

Supplementary Figure 27. Manhattan plot for genes important for the network for separating between cases and controls for atrial fibrillation. Similarly, to coronary artery disease we suspect we need a larger sample size.

##### 4.11 Diabetes

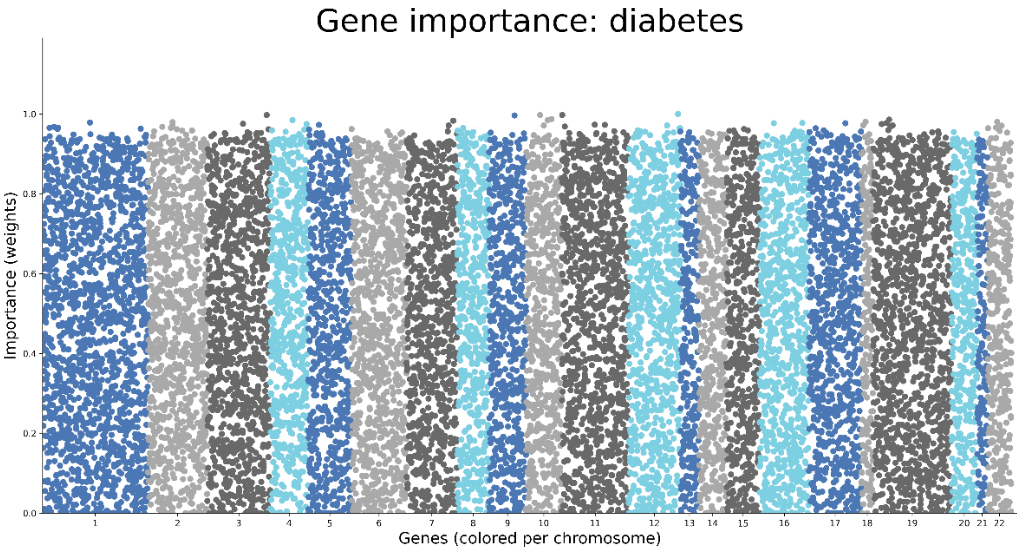

Supplementary Figure 28. Manhattan plot for genes important for the network for separating between cases and controls for diabetes. Although we are underpowered, there seems to be more distinction between genes than for the CAD and AF.

#### Supplementary 5. Rotterdam Study

##### 5.1 Eye color – blue vs rest

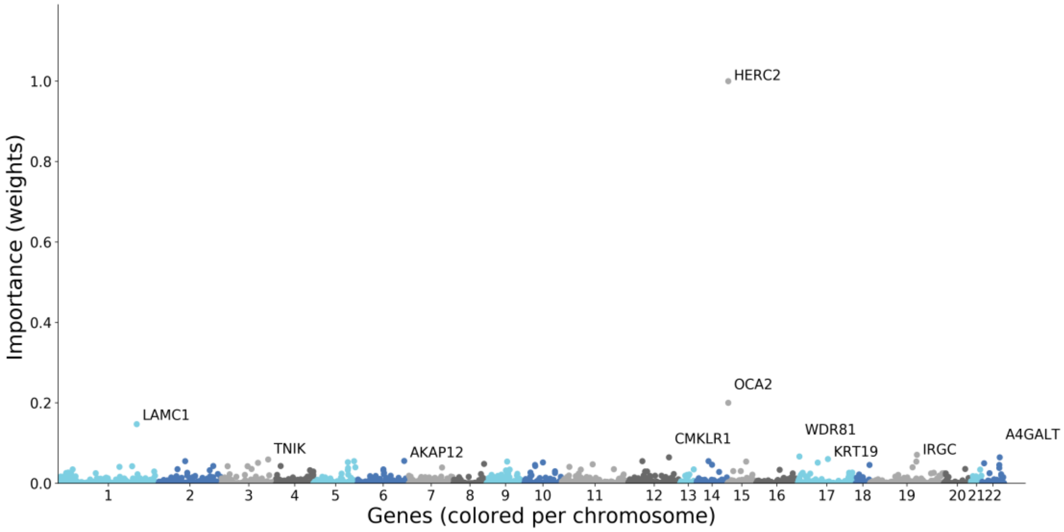

Supplementary Figure 29. Manhattan plot for genes important for the network for separating between blue and other color eyes.

#### Supplementary 6. Regression

For regression tasks, mean squared error is used as a loss function in combination with ReLu activations. Using the UK biobank WES data, the explained variance for height was 31% using linear regression, whereas the network achieved 9% explained variance without any optimization. We expect the network to outperform linear regression with more optimization, since the network has more modelling capabilities than linear regression. In the framework the type of task, regression or classification is automatically determined using the phenotype labels.

**Supplementary 7. Estimating upper bound classification accuracy**

Imagine the following hypothetical situation: for every person in our dataset there exists a monozygotic twin. In our experiment we use the monozygotic twin for the prediction of a phenotype.
Our first phenotype of interest is schizophrenia. It has a concordance rate = 0.5 in monozygotic twins and a prevalence of roughly 1%. We can use these to construct a confusion matrix for our phenotype of interest:

*What if the first twin is schizophrenic?*

- The chance that the twin in our dataset is also diseased is simply the concordance rate, 50%.
- The chance that we misclassify the twin in the real world as a false positive is 1-concordance rate * 100% = 50 %

*What if the first twin is healthy?*

- The chance that the second twin is healthy is higher than the 1-prevalence. Thus, bigger than 99% since the twins share genetically the same code. Let's make this 100% since we are interested in the maximum performance.
- The chance that we misclassify the twin as a false positive is smaller than the prevalence so a maximum of 1%. So let's make this 0% for the best performance.

Resulting in the following confusion matrix:

| | |
|---|---|
| True positive (concordance rate)<br>0.5 * 6245 = **3122** | False Positive (1- concordance rate)<br>(1-0.5) *6245 = 3123 |
| False negative (< prevalence)<br>0* 4969 = **0** | True negative (> (1-prevalence))<br>(1-0) *4969 = **4969** |

Maximum Accuracy = predicted correctly / predicted wrong
= (3122 + 4969) / (4969 + 6245) = 0.72
The maximum accuracy I can reach with this distribution of cases and controls is ~72%.

*This is the perfect classifier if we purely use genetic data. No machine learning or deep learning model can do better without adding (environmental) information.*

The confusion matrix can be used to calculate more metrics such as sensitivity, specificity and $F_1$-score. The sensitivity and specificity can be plotted in the ROC curve to control for overfitting.

#### Supplementary 8. References
